# Quantitative spatial connectomics of liver reveals a partially exposed sympathetic network that integrates multiple cell types across the portal tract

**DOI:** 10.64898/2026.10.26.761888

**Authors:** Luke A. Noon, Giulia Casal, Mario Soriano-Navarro, Anne-Laure Cattin, Rafael Camacho, Carlos Acosta Umanzor, Carmen Rojo Sanz, Arantxa Leal Tassias, Julia Fernandez-Rodriguez, Elisabeth Verboven, Jose Antonio Gomez Sanchez, Marousa Darsinou, Alex Power, Antonella Riccio, Emma R. Andersson, Jemima J. Burden, Alison C. Lloyd

**Author notes:** These authors contributed equally to the manuscript (Luke A. Noon, Giulia Casal).

## Abstract

Peripheral tissues are innervated by sparse axonal networks that regulate homeostasis, regeneration and disease, yet their three-dimensional architecture and cellular targets remain largely unknown. Here, we present an EM workflow for quantitative 3D spatial connectomics to define these networks. We identify the mouse liver connectome as a meshwork, rather than a tree-like structure, of mainly sympathetic axons, in which partial Schwann cell coverage is associated with exposed varicosities along the length of individual axons with large regions of the network devoid of glial coverage. Axon tracing identified axonal terminals throughout the network, but the vast majority of cellular contacts involved exposed “en passant” varicosities. While both terminals and varicosities innervated the vasculature, most synaptic-like contacts involved hepatocytes, fibroblasts, macrophages, and the stem-cell niche, with all individual axons contacting different cell types along their length, highlighting the network nature of liver innervation, the potential for co-ordinating tissue function and demonstrating the power of this approach for decoding connectivity in peripheral tissues.

## Introduction

Peripheral nerves regulate the homeostasis, function, and repair of all peripheral tissues. While studies on the innervation of skin and muscle have identified specific cellular targets such as the neuromuscular junction and sensory structures within the skin and defined the nature of these interactions^1–3^, our understanding of how many peripheral tissues are innervated remains rudimentary. Recent advances in 3D imaging have enabled the tracing of nerves within specific organs, which has begun to shed light on the complexity of tissue innervation^4–9^, but while these studies identify the major neuronal cell types innervating a specific tissue, they lack the resolution to identify the nature of the innervation, such as whether and how specific cell types within a tissue are contacted and the communication network that potentially coordinates tissue function. In the central nervous system (CNS), extensive 3D Electron Microscopy (EM)-connectomics-based mapping has revolutionised our knowledge of brain neuroanatomy, connectivity and complexity^10–15^, however these approaches have not yet been applied to systematically map innervation in peripheral tissues, which is fundamental to our understanding of how the nervous system controls the complex processes involved in tissue physiology.

The ultrastructural characteristics and cellular targets of nerves in the mammalian liver remain largely uncharacterised. Understanding this architecture is important, as liver innervation regulates liver metabolism, inflammation, bile and fluid homeostasis, together with adaptive processes such as lymphatic remodelling, and regeneration after injury, but the mechanisms remain obscure^16,17^. In both humans and mice, liver innervation is almost exclusively composed of sympathetic axons that enter along the hepatic artery and the portal vein and track along these vessels as they branch along the portal tract^18–20^. The major cell types in the liver express adrenergic receptors and consistent with this, dysfunction of the sympathetic network changes liver metabolism and is associated with the development of metabolic disorders such as metabolic-dysfunction-associated fatty liver disease^21,22^. Interestingly, the sympathetic fibres are acutely sensitive to metabolic stresses, such as obesity, resulting in dramatic neuropathies, with loss of much of the terminal network. However, the network is restored upon resolution of the metabolic stress demonstrating the regenerative capabilities of the sympathetic nerves which innervate the liver and the potential for manipulating neuronal signalling for therapeutic purposes^18,19,22^.

The peripheral neural networks that coordinate the function of tissues differ from the brain, in that far sparser networks relay the signals that co-ordinate behaviours within individual tissues, compared to the highly dense neural networks that process information in the brain. Mapping of these networks therefore requires an approach that can first identify the nature and location of the sparser neuronal networks, coupled with ultrastructural analyses that can determine how the networks interact with individual cell types and structures within a specific tissue. To achieve this, we have developed a cross-scale connectomics workflow that combines nerve targeting using Correlative Light and Electron Microscopy (CLEM), together with whole-tissue imaging and cross-scale volume electron microscopy (vEM), enabling large-scale 3D reconstruction of the peripheral nerve microenvironment at nanometre resolution. Applied to the liver, across multiple animals, this quantitative approach has identified a network that challenges our perceptions of how neural pathways control tissue function. Instead of a tree-like branching structure that terminates at individual structures, we find an intricate meshwork of sympathetic axons organised across the portal tract. Within this complex, bi-directional network, most cellular contacts occur “en passant” along the length of individual axons, with most interactions involving non-vascular cell types such as hepatocytes and fibroblasts. In addition, large tracts of the network are devoid of Schwann cell coverage, countering the view that peripheral axons are always ensheathed by glia along their lengths^23,24^. Tracing of individual axons identified multiple points of contacts, often with different cell types, providing a strategy to potentially co-ordinate tissue function. Together this work provides a transferable, quantitative workflow to analyse peripheral connectomes, and identifies the complex network of exposed sympathetic axons that innervate cell types across and along the portal tract of the liver.

## Results

### The portal tract is innervated by a meshwork of sympathetic neurons

Autonomic nerve bundles enter the liver alongside the hepatic artery (HA) and extend throughout the connective tissue of the portal tract, where they run adjacent to the portal vein (PV), bile ducts, lymphatic vessels and the hepatocytes that surround the portal space^18,19^ (Figure 1A). In order to understand how these nerve bundles branch to subsequently innervate the liver, we took a cross-scale approach; combining whole tissue imaging and ultrastructural analysis - using vEM to map nerve connectivity throughout the organ. Compared to the CNS, peripheral nerves are more sparsely and non-uniformly distributed throughout tissues, making it more challenging to map neural networks in peripheral tissues. To circumvent this issue, we used transgenic mice in which enhanced green fluorescent protein (eGFP) is robustly expressed in Schwann cells (SCs) *(Plp-eGFP* mice)^25,26^. SCs, the glial partners of peripheral neurons, have a larger cytoplasmic volume compared to ultrafine axons, which allows easier identification of the neural network by CLEM for subsequent ultrastructure analysis. In addition, the use of endogenous fluorescence negated the requirement for permeabilisation, which better preserved the ultrastructure of the tissue, providing a scalable, glia-based strategy for the cross-scale characterisation of peripheral nerve architecture.

**Figure 1:**
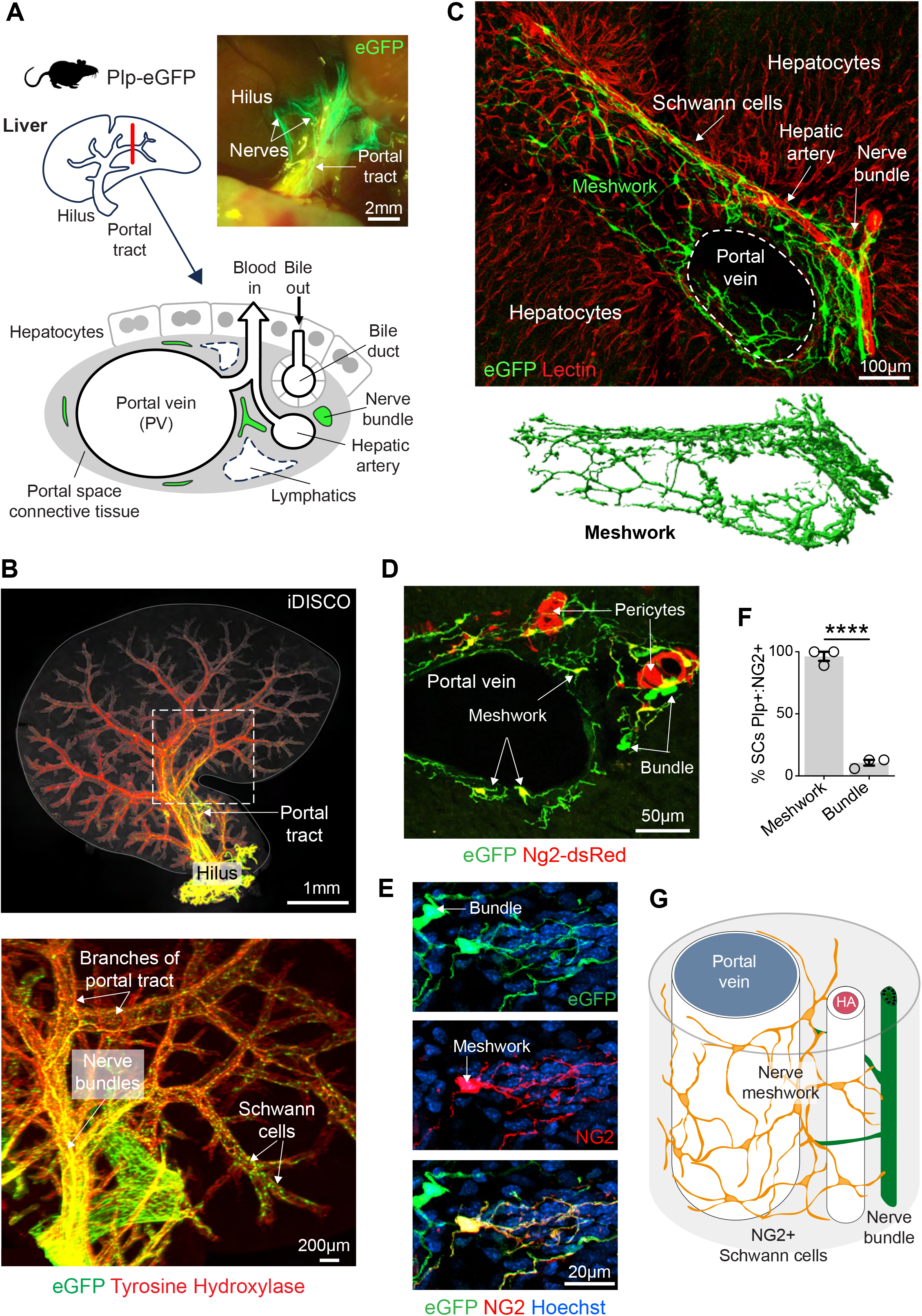
The portal tract is innervated by a meshwork of sympathetic neurons. (A) Schematic of the liver portal tract. Image shows a fluorescent wide-field image of the liver showing nerve bundles (labelled by SC expression of eGFP (green)) entering the base (hilus) of the liver via the portal tract. (B) Representative light-sheet images of an iDISCO-cleared caudate lobe from Plp-eGFP mice immunolabelled to detect sympathetic axons (Tyrosine hydroxylase, red) and SCs, (eGFP, green) (see MovieS1). (C) Representative confocal image of a CLARITY-cleared liver labelled to detect blood vessels (lectin Cy3, red) and SCs (eGFP, green). Lower image shows an Imaris 3D reconstruction of SCs illustrating their meshwork organisation. (D) Representative confocal image of a transverse section of the portal tract of a liver from Plp-eGFP:Ng2-dsRed mice to detect SCs (eGFP,green) and NG2-expressing cells (ds-Red, red). (E) Representative confocal images of the portal tract of mouse liver immunolabelled to detect NG2 (red) and SCs (eGFP, green). Only SCs within the meshwork express NG2. (F) Quantification of (E) showing the percentage of NG2+ SCs (eGFP+) in the meshwork or in larger bundles (n=mice, mean ± SEM). (G) Schematic to illustrate the sympathetic network across the portal tract. HA = hepatic artery. ****p<0.0001 by unpaired t-test.

Initially, using widefield fluorescence imaging of intact livers from *Plp-eGFP* mice, we observed large eGFP+ nerve bundles entering the portal tract at the base (hilus) of the liver, as expected (Figure 1A). To visualise their trajectory throughout the liver, we performed whole-organ clearing followed by light sheet microscopy. Co-staining for tyrosine hydroxylase (TH), a marker for sympathetic neurons, revealed the extent of the sympathetic nerve network that extended along the entire length of the portal tract (Figure 1B, Movie S1) with associated SCs visible across the network. Consistent with previous findings that the liver is primarily innervated by sympathetic neurons^18,19^, we co-stained sections of the liver for TH and the universal axonal marker PGP9.5 and found that within the portal tract the vast majority of the axons (>97%) were sympathetic (Figures S1A-B).

To characterise how this neural network aligned with the portal vasculature, we performed confocal imaging of tissue-cleared livers from *Plp-eGFP* mice injected with fluorescent lectin to label blood vessels. In line with previous findings, this revealed SC-labelled nerve bundles aligned along the hepatic artery^4,18,19^. The bundles then branched into a mesh-like structure that extended across and along the length of the portal tract (Figure 1C), a finding supported by 3D nearest-neighbour analysis of SC distribution (Figures S1D-E).

Terminal (“perisynaptic”) SCs at neuromuscular junctions express NG2, a universal pericyte marker, whereas both myelinating SCs (mSCs) and non-myelinating SCs do not ^27,28^. To identify potential terminal SCs within the sympathetic network, *Plp-eGFP* mice were crossed to *Ng2-dsRED* mice^29^. Confocal analysis of cross-sections across branches of the portal tract showed a meshwork of SCs, spreading across the portal space branching away from the large nerve fibres around the hepatic artery (Figure 1D). As expected, SCs in the larger nerve bundles along-side the hepatic artery, did not express *NG2*-directed dsRed (Figure 1D). In contrast, the vast majority of SCs within the meshwork expressed dsRed, suggesting a distinct, more terminal function (Figure 1D). This finding was confirmed by immunostaining for NG2 (Figure 1E-F), which because of its plasma membrane location, further highlighted the distinct morphologies of the SCs within the network: SCs within bundles displayed a conventional Remak SC morphology, whereas NG2+ SCs within the meshwork had a highly distinctive multipolar and ramified appearance (Figures 1E and S1F-G). Imaris 3D reconstruction confirmed that the vast majority of SCs within the network were NG2+ (Figure S1G). Together these findings show that the liver is innervated along the portal tract by a meshwork of sympathetic axons, associated with NG2+ SCs, indicative of extensive interconnections throughout the portal tract (Figure 1G).

### Volume-EM reveals a complex axonal network within the portal tract

While 3D confocal imaging allowed us to visualise the extensive sympathetic axonal network along and across the portal tract, it lacked the resolution to determine the nature of the network and how it innervates cells of the liver. VEM is the gold-standard technique for connectomics, supplying the 3D resolution and volume required to analyse the architecture and trajectory of nerve fibres and their sites of interactions, and has been used extensively and successfully to analyse neural networks in the brain^12,14,15,30^. In the PNS however, the challenge of finding neurons in tissue with a sparser axonal network, required the development of a new workflow. We established a volume CLEM (vCLEM) protocol, that used eGFP+ SCs to identify axonal networks for targeting, tracking and 3D reconstruction (Figure S2A and Movie S2). This workflow optimised the targeting of high-resolution image acquisition by combining CLEM guided, hierarchical imaging of serial section arrays, using non-destructive, automated, Array Tomography Scanning Electron Microscopy (AT-SEM) (see Methods). This approach enabled the efficient use of SEM resources to acquire only the volumes of liver that contained neurons, that could then be reconstructed with high fidelity (5-10 x 70nm voxel size). Analysis workflows incorporated cloud-based vEM data sharing and team-based annotation, in association with Webknossos ^31^, accelerating throughput and ensuring reproducibility. Importantly, the increased efficiency in targeting permitted a more high-throughput approach allowing the quantification of structures from multiple animals.

In total, 12 interactive vEM datasets (Table S1) have been deposited online, representing a cumulative total of 815,851μm^3^ of liver tissue across 3725 serial sections from 4 animals. These are publicly available allowing community analysis of these information-rich datasets. Analysis of skeletonised axons revealed a range in axonal density across the portal tract from as low as 0.006μm/μm^3^ to more dense regions (0.137μm/μm^3^) (Table S1), which despite targeting by CLEM, at their densest are still ∼60 fold lower than those reported in the CNS (7-9μm/μm^3^)^32^. This underlines the need for specialised workflows to analyse the sparse and unevenly distributed peripheral neural networks.

Analysis of axonal trajectories within the volume datasets identified an interconnected network of axons consisting of larger (>10 axons) and smaller bundles together with areas rich in single axons, consistent with the network architecture visualised by light microscopy (Figures 1C and 2A and Table S1). However, the higher resolution of vEM allowed the visualisation of the trajectories and interactions of individual axons permitting us to characterise the nature of liver innervation. In a typical example shown in Figure 2A, we identified a total of 114 axon skeletons in 1.6 x10^5^ μm^3^ of tissue, within the portal space. 49% of the axons in this dataset (56) were within 3 nerve bundles (>10 axons), found close to the hepatic artery or lymphatics, in line with observations from the confocal analysis (Figures 1C-D). The remaining 51% of axons (58) formed an interconnected meshwork of branching axons, either in small bundles or single axons, running across the portal tract, with most single axons found around the hepatocyte border (Figure 2A and S2B), a finding replicated across the other datasets (Table S1). Within this network, 12/114 (∼10%) of axons terminated within the volume, suggesting that axons were innervating regions across the portal tract.

**Figure 2:**
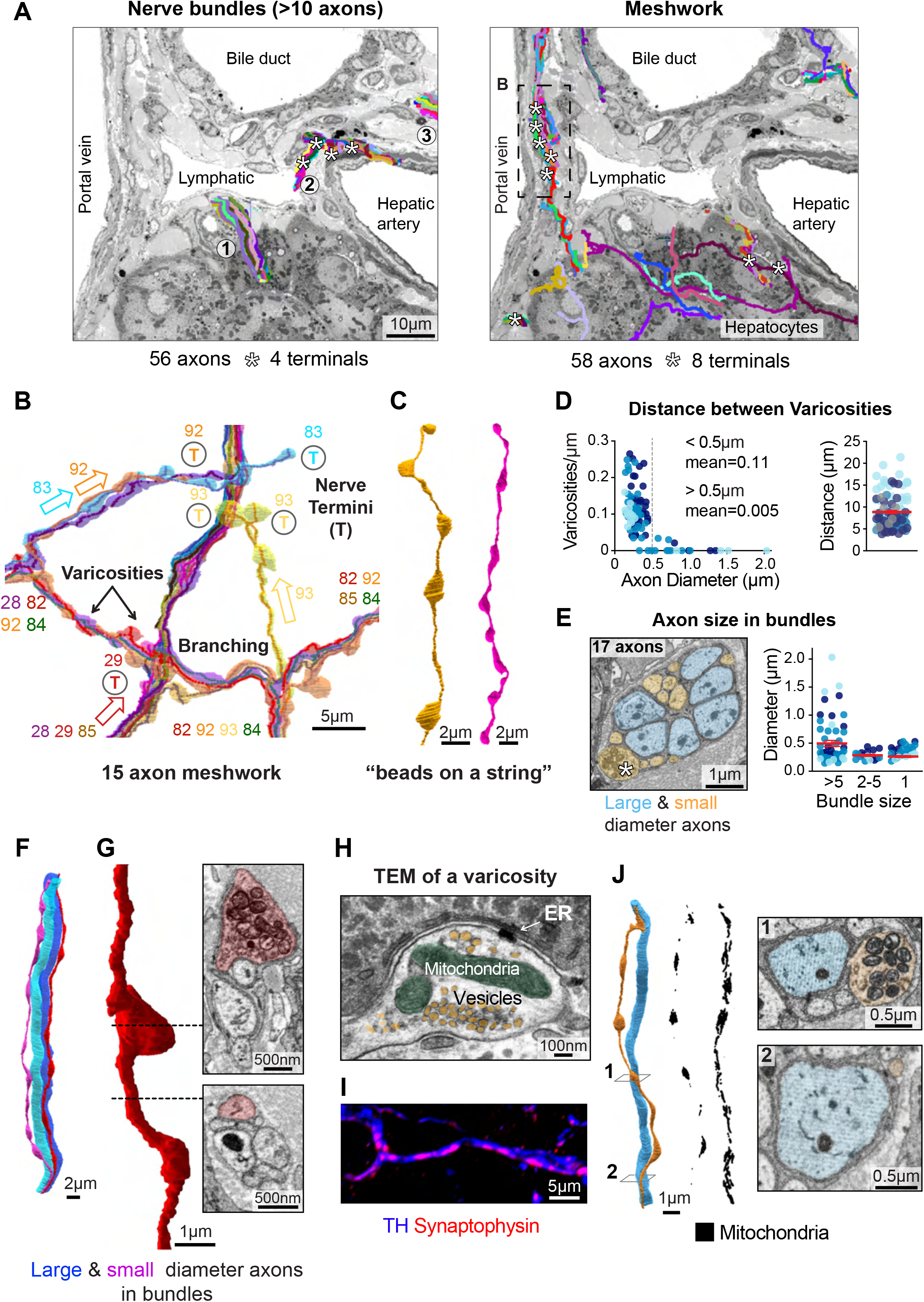
Volume EM reveals a complex axonal network within the portal tract. (A) Images show the 3D reconstruction of axons within a 1.6 x10^5^ µm^3^ volume of portal tract tissue acquired using AT-SEM. The 117 axons within the dataset were skeletonised and superimposed onto an EM section to indicate their spatial distribution. The left image shows the 56 axons present in 3 nerve bundles. The right image shows the 61 axons that form a meshwork within the portal tract. Axon terminals indicated by white stars. (B) Segmented reconstruction of a subgroup of 15 axons from the dashed box in (A). Arrows indicate axon directionality inferred from terminal locations (T). Axons are numbered and colour coded as the online dataset. (C) Examples of segmented small diameter axons showing periodic varicosities along their length. (D) Quantification of varicosity density dependency on axon diameter and shows that only axons with a diameter of <0.5µm have multiple varicosities. Each dot represents an individual axon (n=3 mice). Right: Calculated distance between varicosities, each dot represents an individual axon (n=3 mice). (E) EM image of a bundle containing a mixture of large and small diameter axons. White star indicates a varicosity of a small diameter axon within the bundle. Graph shows quantification of the size of axons within small and large bundles and shows that many small axons are present in bundles. Each dot represents an individual axon in one of 3 mice (shades of blue). Data shown as mean ± SEM. (F) Selected segmented large and small diameter axons from (E). Note varicosities are only observed in the small diameter axons. (G) Segmented 3D reconstruction of an axon with EM images showing cross-sections at and after a varicosity. (H) Representative TEM image of a varicosity, pseudo-coloured to show mitochondria (green) and vesicles (mustard). The aligned ER of the contacting cell is indicated with an arrow. (I) Representative confocal image of mouse liver immunolabelled to detect synaptophysin (red) and sympathetic axons (TH, blue). (J) Segmented reconstruction of a large (blue) and small (orange) diameter axon within a bundle. Segmented mitochondria (black) within each axon are shown. EM images of the indicated slices showing (1) a varicosity and (2) a region between the varicosities.

To further analyse the meshwork, we initially focused on a subgroup of 15 axons within this dataset (total length 0.95mm) within a meshwork-like structure running in parallel to the portal vein (Figures 2A-B). This region contained small bundles, (ranging in size from 2-10 axons), together with a single axon. 3D segmentation of the axons revealed several interesting features: Firstly, 4 of the 15 axons terminated within this region, with some terminating within small bundles (Figures 2B and S2C and Table S2). Secondly, analysis of the trajectories of axons that terminated within the dataset (and hence revealing their directionality), showed that within bundles, some axons were running in an antiparallel manner (Figure S2D), a finding confirmed in other datasets. Thirdly we observed terminals converging from opposing directions within small bundles (Figure S2E). Together these analyses highlight the 3D complexity of the neural network that innervates the portal tract of the liver.

A further observation was that prominent varicosities could be observed along the length of axons within the network (Figures 2B-C). These axonal bulges, often referred to as “beads on a string” have been speculated to be associated with “en passant” nerve signalling^33^, suggesting that autonomic nerves in the liver signal along their length, rather than solely at their terminals. Analysis of individual axons from across our datasets showed that varicosities were ubiquitous in small calibre axons (diameter <0.5μm) but were mostly absent from the larger diameter fibres, found within bundles (>0.5μm diameter) (Figure 2D). Analysis of the frequency of the varicosities found an average distance of ∼9μm between varicosities (Figure 2D), which is similar to the linear density of synapses along axons within the CNS, that are associated with network signalling ^32^. Remarkably, but consistent with the meshwork nature of the network, we observed that the nerve bundles did not only consist of larger axons but also many smaller axons (Figure 2E). These smaller axons also had prominent varicosities showing that the presence of varicosities is a property of the size of an axon rather than its location within the network (Figures 2E-F).

Consistent with a role of signalling at varicosities, these regions were often packed with both mitochondria (Figures 2G-H) and secretory-like vesicles, as those in the CNS (Figure 2H). Moreover, immunofluorescence staining for synaptophysin, a marker of synaptic vesicles, identified similar structures along the sympathetic axons that mirrored the location of varicosities along the length of individual axons (Figure 2I). This was in contrast to the larger axons which showed a consistent density of mitochondria along their length (Figures 2J).

To determine whether periodic varicosities are a property of sympathetic axons outside of the liver, we cultured rat sympathetic axons and analysed their structure by both light microscopy and EM. Interestingly, both analyses confirmed the existence of varicosities along their length with a similar spacing (5μm), together with an accumulation of mitochondria and vesicles within the varicosities, which stained positively for synaptophysin, indicating they may be primed for signalling (Figures S2F-H). Further analysis of the positioning of the varicosities along the length of the axonal skeletons from separate datasets showed that varicosities from different axons, particularly those in small bundles, were frequently found in clusters, rather than being distributed randomly within the region (Figure S2I-K). This further suggests co-ordination of signalling within the axonal network and emphasises the potential interconnectivity of the portal network.

### Axonal terminals within the network are frequently free of Schwann cell coverage

Peripheral innervation is arguably best characterised in muscle and skin^34–36^. In both cases, axonal terminals mostly innervate specific cell types or structures-such as muscle cells or blood vessels; or sensory structures, such as Meissner corpuscles. In this context, they act to either control functions such as muscle or vessel contraction or relay information related to the sensing of the environment^2,37^. Dependent on these roles the terminals have highly specialised endings that enable their function^3,9,34^. Schwann cells have been thought to cover the full-length of peripheral axons providing essential metabolic and protective support for these exceptionally long structures with tissue-specific, often highly specialised, terminal SCs that enwrap the axonal terminals and can participate in signalling to and from the structures they innervate^3,23,24,38–42^. Described exceptions are rare but include partially SC-free nociceptive sensory nerve endings within the epidermis of the skin ^40^ and sensory nerve endings within the corneal epithelium^43^.

Analysis of our datasets identified a total of 54 axon terminals (from n=4 mice) within the portal networks (Table S2). These could be grouped into four categories based on firstly, the structure of the axon terminal as either blunt-ended or bulbous: Blunt terminals (16/54) resembled the trunk of the corresponding axon in terms of width and a similar low density of intracellular structures but tapered to an ending (Figure 3A-B). In contrast, the more common bulbous terminals (38/54) were larger and contained more visible organelles such as mitochondria and numerous vesicles of varying sizes, indicative of signalling termini (Figures 3A and 3C)^44^. The second categorisation was whether the terminals were associated with a terminal SC. Somewhat surprisingly, the majority (70%) of both blunt and bulbous axonal terminals were not associated with a SC (Figures 3A-B). Further analysis of the SC-free terminals indicated that the axons were frequently devoid of SC coverage for substantial lengths from their terminals (Figures 3D-E). This analysis included axons for which we could follow the axon to the first encountered SC, which could extend >60μm (Figure 3E). Another group extended even further, up to >90μm, and beyond the volume of the datasets, indicating the axons were free of SCs for considerable distances (Figure 3F-G).

**Figure 3:**
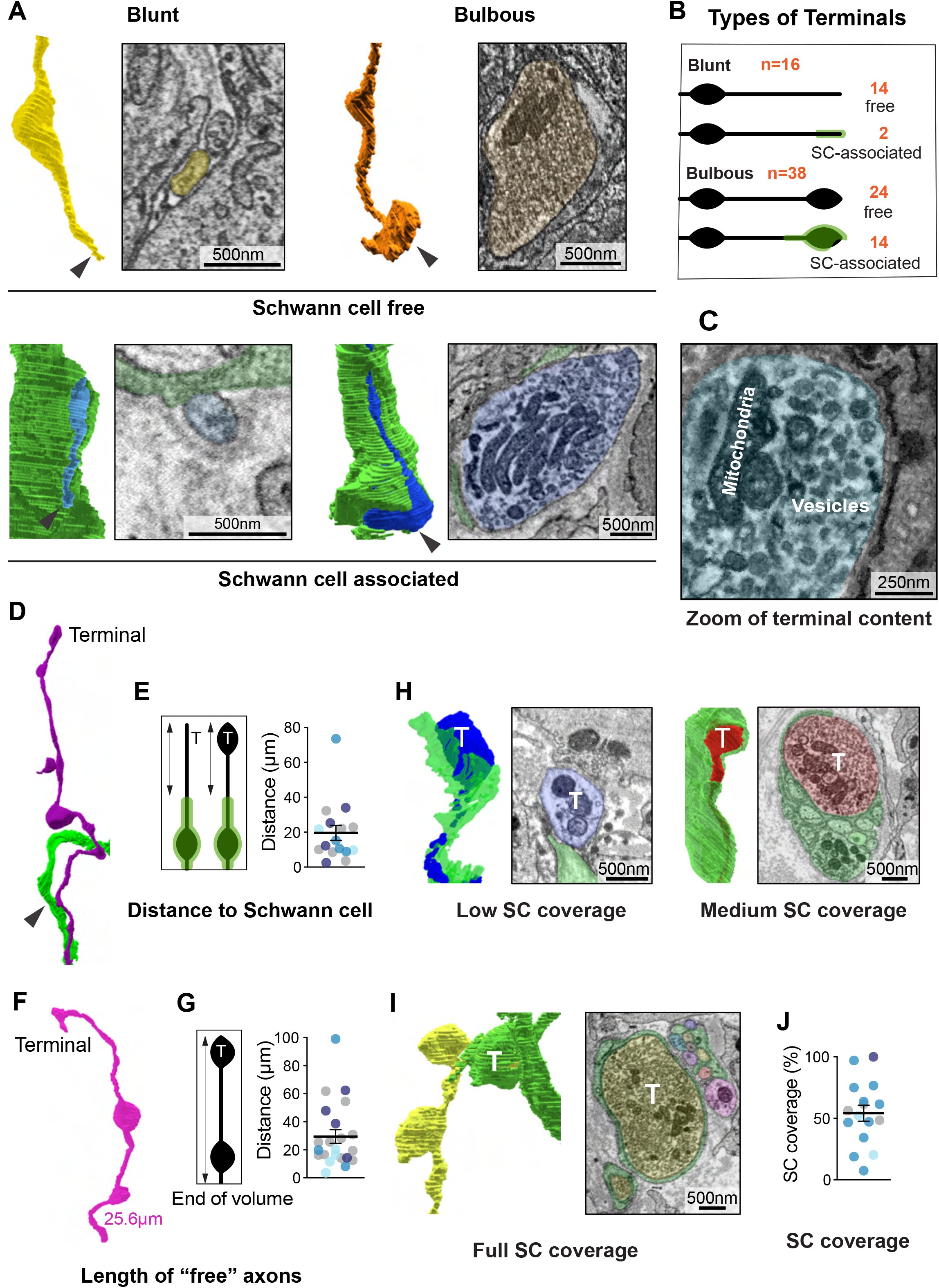
Axonal terminals within the network are frequently free of Schwann cell coverage. (A) Representative examples of blunt and bulbous axon terminals. Segmented axons and associated SCs are shown, together with an EM image of the terminal region, pseudo-coloured to indicate axons (yellow/blue) and SCs (green). Black arrowheads indicate the terminals. (B) Schematic illustrating the two terminal morphologies identified (blunt and bulbous) and their association with SCs (green) with the numbers of each indicated (see Table S2). (C) Representative EM image of a bulbous terminal showing the terminus (pseudo-coloured blue) is packed with mitochondria and vesicles. (D) Example of a segmented axon (purple) illustrating the distance from the terminal to the nearest SC (green) contact. (E) Graph shows quantification of the distance from each terminal to the first SC contact. Each point represents a terminal from n=3 mice; data is shown as mean ± SEM. (F) Example of a segmented terminal showing the distance from the terminal to the end of dataset without SC coverage. (G) Graph shows quantification of the SC-free distance from axon terminal to the end of the dataset. N=3 mice, mean ± SEM. (H) Examples of a terminal showing a low and medium degree of SC coverage. Images show the segmented axonal terminals (blue/red) and associated SCs (green) with associated EM images. T=terminal. (I) Example of a terminal completely covered by a SC. Image shows the segmented axonal terminal (yellow) and associated SC (green) with associated EM image. (J) Graph shows quantification of the degree of SC coverage of terminals associated with a SC. N=3 mice (different colours), mean ± SEM.

Terminals that were associated with a SC, showed a variation in the degree of SC coverage, with some showing minimal coverage and most having at least some free surface that could potentially allow more direct signalling to target cells (Figures 3H and 3J). Interestingly, we found one example in which a terminal was fully covered by a SC, suggesting that not all axonal terminals signal directly to other cell types within the liver (Figure 3I-J).

### The axonal network of the portal tract is only partially covered by Schwann cells

As might be expected, axons within the larger bundles (>10 axons) were enveloped by non-myelinating SCs in structures resembling classic Remak bundles (Figure S3A). However, across the meshwork regions, we observed a wide variation in SC coverage of axons, ranging from fully ensheathed axons within some smaller bundles (2-10 axons) to largely exposed single axons (Figure 4A and S3B-C). However, more remarkably we observed regions of the network with no detectable SC coverage (Figure 4A-B and S3D). These SC-free meshwork regions were most prevalent away from the vessels and closer to hepatocytes, with others located within the stromal spaces between the vessels (Figure 4A). In particular, the mostly SC-free, single axonal network bordering the hepatic border, was extensive, forming a mesh interfacing between the hepatocytes and stromal space (Figure 4B and S3D-E and Movie S3). To confirm the somewhat surprising finding that large tracts of the axonal network were free of SCs, we used confocal and Imaris analysis of livers from *Plp-eGFP* mice labelled for TH and observed substantial tracts of sympathetic axons not enveloped by SCs that tended to be in regions apposing the hepatocyte plate (Figure 4C and quantified in S3F).

**Figure 4:**
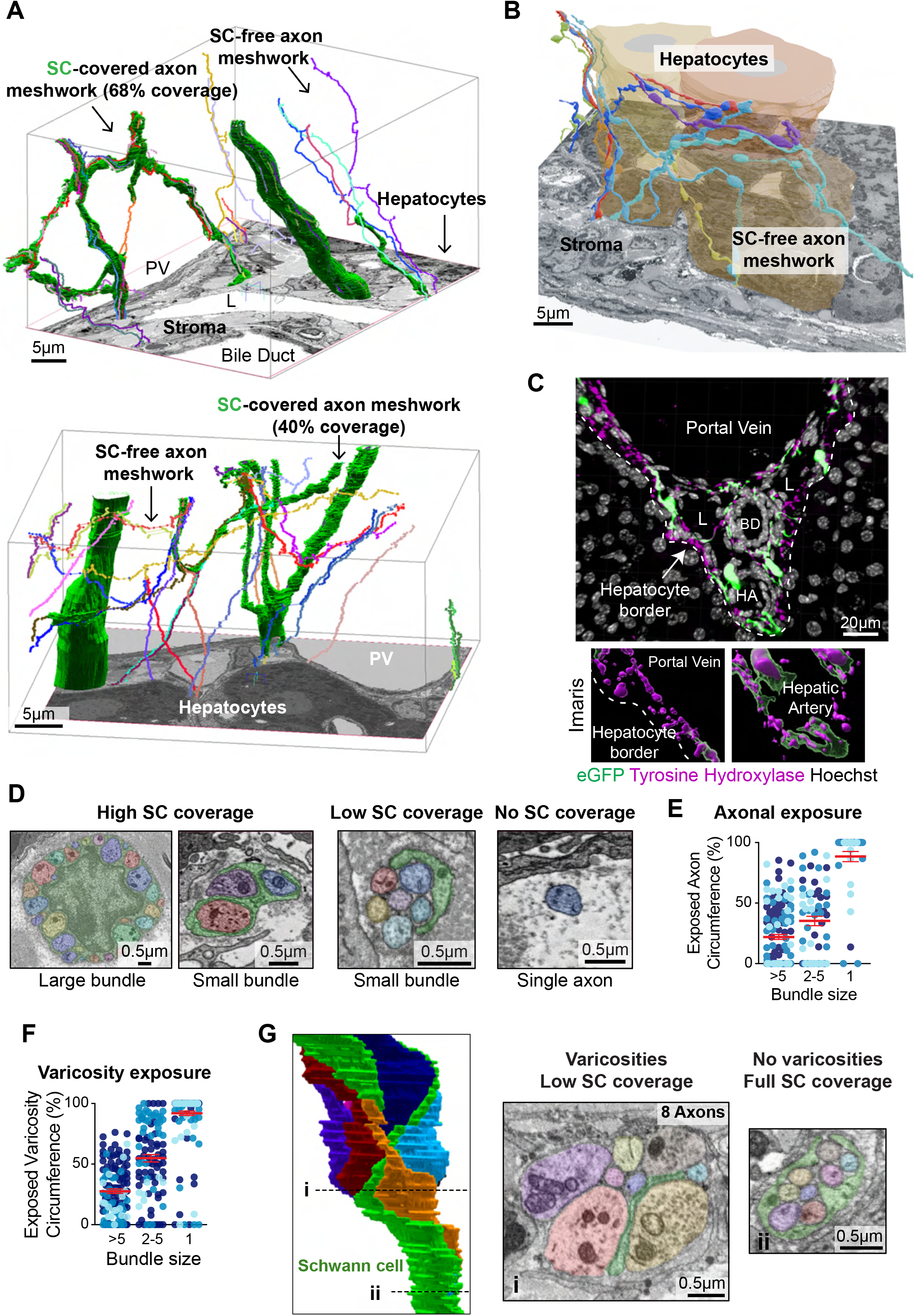
The axonal network of the portal tract is only partially covered by Schwann cells. (A) 3D reconstructions of the portal tract showing axonal bundles and regions of the axonal meshwork with SC coverage or free of SCs. Images show skeletonised axons together with segmented SCs (green), superimposed on an EM section to localise the network relative to the portal vein (PV), lymphatics (L) bile ducts and hepatocyte border. (B) Representative example of a region of the axonal network that is free of SC association, superimposed on an associated EM section that shows a meshwork of segmented, SC-free axons abutting the hepatocyte border (see Movie S3). (C) Representative confocal image of the portal tract immunostained to detect sympathetic axons (TH, purple) and SCs (eGFP, green). Lower images show Imaris reconstructions (HA), highlighting reduced SC coverage in the hepatocyte region. (D) EM images showing variability in the degree of SC coverage across nerve bundles of different sizes. (E) Graph shows quantification of the percentage of exposed axon circumference relative to bundle size. Each point represents a single axon from n=3 mice (different blue shades), >5 (131 axons), 2-5 (65 axons), single (49 axons) analysed. Red lines show mean ± SEM. (F) Graph shows quantification of the percentage of exposed varicosity circumference relative to bundle size. Each point represents a varicosity from n=3 mice (different blue shades), >5 (125 varicosities), 2-5 (97 varicosities), single (171 varicosities) analysed. Red lines show mean ± SEM. (G) Segmented 3D reconstruction (left) and EM images (right) illustrating partial SC coverage of axons within a nerve bundle of 8 axons, highlighting (i) a cluster of exposed varicosities and (ii) a region with full SC coverage.

Analysis across datasets confirmed that axonal exposure increased progressively with decreasing bundle size (Figures 4D-E). Axons within larger bundles (>5 axons) were all associated with SCs but even within some of these bundles there were partially exposed axons, although they mostly remained within the basal lamina of the associated SC (Figure 4D-E and S3G-H). In contrast, smaller bundles (2-5 axons) showed greater variability of SC coverage, with some axons fully covered, whilst others showed only very minimal coverage (Figures 4D-E). Remarkably, single axons tended to be uncovered, suggesting that once axons leave a SC bundle, they are often no longer associated with SCs (Figures 4D-E). As shown in the previous section, some exposed axons were uncovered from the terminals, however we also frequently observed mostly SC-free sections of axon flanked by SC interactions (Figure S3I-J) resulting in exposed internal regions of up to 35μm between sequential SC contacts (Figures S3J-K).

The finding that the meshwork of sympathetic axons permeating the portal tract is only partially covered by SCs has potential implications for how the network signals to the cells of the liver, as exposed axons would more readily permit interactions of the varicosities with other cell types. Consistent with this, single axons with prominent varicosities tended to be free of SC coverage (Figures 4B, 4E and S3C) and staining for synaptophysin showed that these regions were rich in synaptophysin+ puncta indicative of sites of potential signalling (Figure S3J). Somewhat more surprisingly, we also found that the vast majority of varicosities within both the smaller and large bundles were also at least partially exposed, which potentially would also permit direct contact of varicosities within bundles with cells of the liver (Figures 4F-G).

### The sympathetic network terminals contact distinct cell types within the liver

To understand how hepatic nerves innervate their local microenvironment, we first mapped the contacts formed between the axon terminals and cells within the liver (Table S2). The different cell types could be readily distinguished by their morphologies as described in Figures S5A-B and Methods. A few terminals interacted with more than one cell type, resulting in more detected interactions than terminals (Figure 5A). As might be predicted, a proportion (28%) of axonal termini were associated with the vasculature including the hepatic artery, portal vein and smaller vessels around the bile ducts (Figure 5A). However, the majority of interactions were with other cell types, with the most common contacts being with fibroblasts (44%), hepatocytes (24%) and macrophages (3.6%) (Figure 5A).

**Figure 5:**
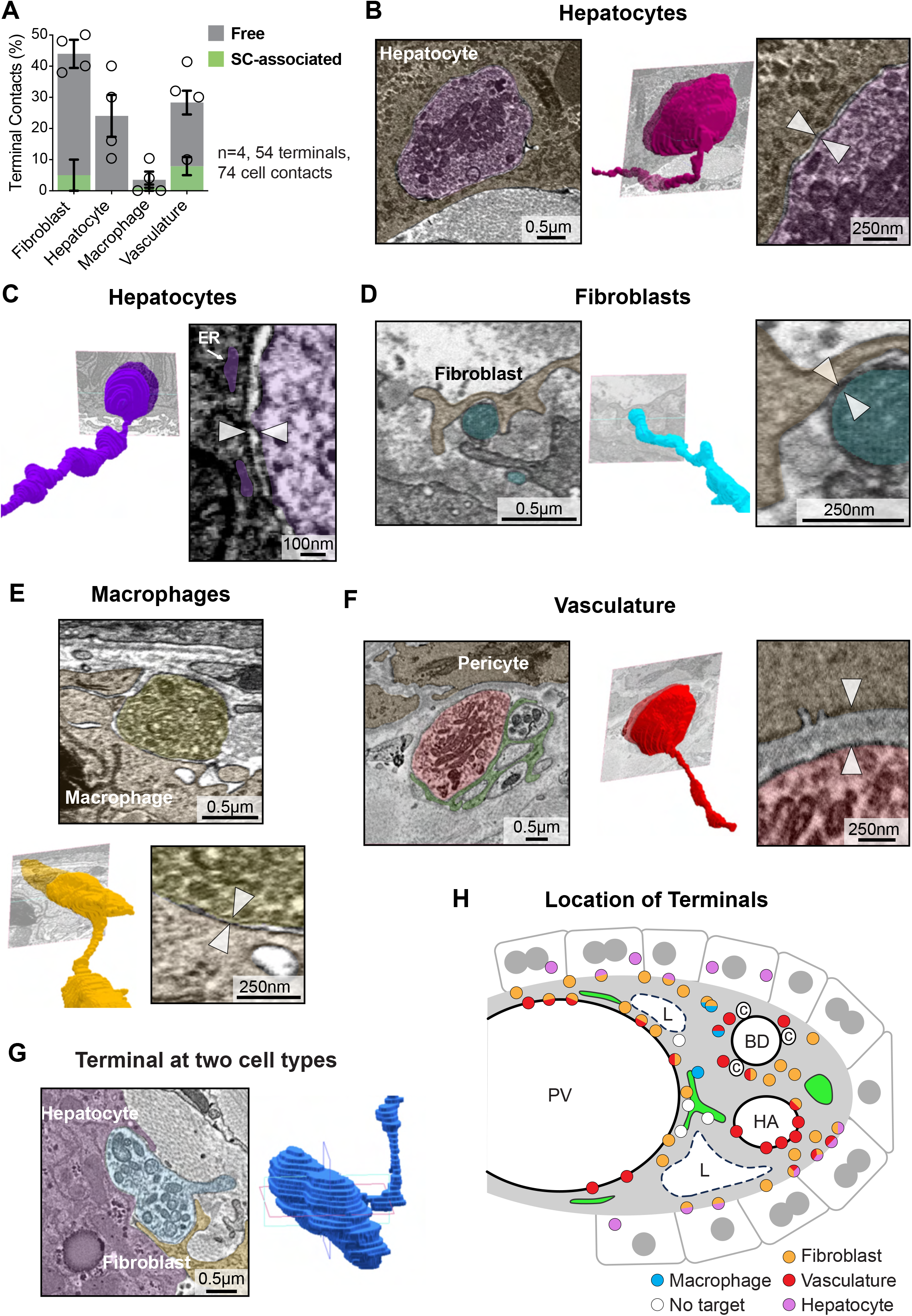
Sympathetic network terminals contact distinct cell types. (A) Graph shows quantification of the indicated cell-types that interact with terminals (Table S2). Terminals are defined as either SC-associated (green) or free of SC coverage (grey). N=4 mice. Data is shown as mean (circles for each animal) ± SEM for free (grey) and SC-associated terminals (green) bars. (B) Representative EM image of a terminal’s interaction with a hepatocyte, together with a segmented volume of the terminal, and a zoomed EM showing the separation between the terminal and the hepatocyte. (C) EM image showing the “synaptic” space (∼20nm) between a terminal and the associated hepatocyte. Arrowheads indicate the synaptic space. Arrow indicates aligned ER within the hepatocyte. (D-F) As (B) showing terminal interactions (pseudo-coloured) with (D) a fibroblast (E) a macrophage and (F) the vasculature. (G) EM image shows a terminal that interacts with two cell-types together with the fully segmented terminal image and is pseudo-coloured to show the terminal (blue), hepatocyte (purple) and fibroblast (yellow). (H) Schematic illustrating the spatial distribution of axon terminals across the portal tract within the analysed datasets.

Interestingly, we observed differences in the nature of the interactions of the terminals with the different cell types. Hepatocyte interactions involved bulbous terminals, packed with vesicles and mitochondria, that frequently protruded into the hepatocyte. Notably, all were free of SC coverage (Figures 5A-C and Table S2). Consistent with a synaptic-like interaction, a distinct and uniform ∼20nm gap could be observed between the terminal and the surrounding hepatocyte^44,45^. Moreover, the endoplasmic reticulum (ER) of the hepatocyte appeared aligned to the contact site, indicative of a signalling interaction (Figures 5C, S4C and Table S2)^46^. In contrast, the interactions with fibroblasts were also direct and involved a similar sized gap, the terminals did not protrude into the cell and there was less visible polarisation at the contact site, with many of the contacts made at thin fibroblast protrusions (Figures 5D, S4C and Table S2). The rarer macrophage interactions were again different, in that the terminals appeared smaller, with fewer vesicular structures and no ER alignment within the macrophages was observed (Figures 5E, S4C and Table S2).

The nature of the interactions with the vasculature was also distinct, in that the majority of these terminals made contact with the basal lamina of the vessels, rather than directly with the cells of the vessels and more were associated with SCs (Figures 5A, 5F and S4C), consistent with previously reported sympathetic interactions with vessels in other tissues^47^. However, in 6/20 interactions, we observed that the terminals penetrated the basal lamina to make direct contacts with pericytes surrounding the vessels (Figure S4D).

While the majority (69%) of terminals innervated a single cell type, some made contacts with more than one cell type. In 15 of the 17 examples, this involved a fibroblast together with another cell type (Figure 5G). Notably a few axons (4) did not appear to terminate at a cell but instead either ended within a bundle or terminated within the extracellular space (Figure S4E). Analysis of the location of all terminals showed they were distributed across the portal tract, rather than to specific regions (Figure 5H) consistent with an interconnective network that coordinates sympathetic signalling across the liver.

### The sympathetic network makes “en passant” connections with multiple cell types

Whilst terminals of the sympathetic network make direct contacts with cells across the portal tract, the vast majority (>97%) of cellular interactions instead occurred at varicosities found along the lengths of the small diameter axons, with nearly all varicosities making direct contact with another cell that was not a SC (Figure 6A and Table S2). In the SC-free regions of the network, mostly consisting of single axons, and frequently found in regions abutting the hepatocytes (Figures 4A-C), close to 100% of varicosities made direct contact with another cell, indicating that all varicosities in these regions may act as signalling nodes (Figure 6A and Table S2). Similarly, varicosities in regions of the meshwork, which contain more small bundles and partial SC coverage mostly made direct contact with other surrounding cells (77.3%). Those that did not were usually associated with a SC protrusion (81.6%), that often appeared to form a signalling contact, based on the alignment of structures within the SC at the contact sites (Figure S5A-C). This finding would be in keeping with the reported nurturing role of glia in supporting peripheral axons^39,48^. Perhaps more surprisingly, close to 50% of varicosities in the small-diameter axons within the larger nerve bundles also made direct contact with another cell type, with partial SC coverage of the varicosities and the orientation of the bundles allowing contact to cells in the parenchyma (Figures 6A and S5D). Some of the exposed varicosities within the bundles did not directly contact a cell but were instead exposed to the extracellular matrix (Figure S5D). Together this indicates that varicosities are interacting with multiple cell types across the network and that even within larger nerve bundles, the sympathetic network may be directly signalling to cells of the liver.

**Figure 6:**
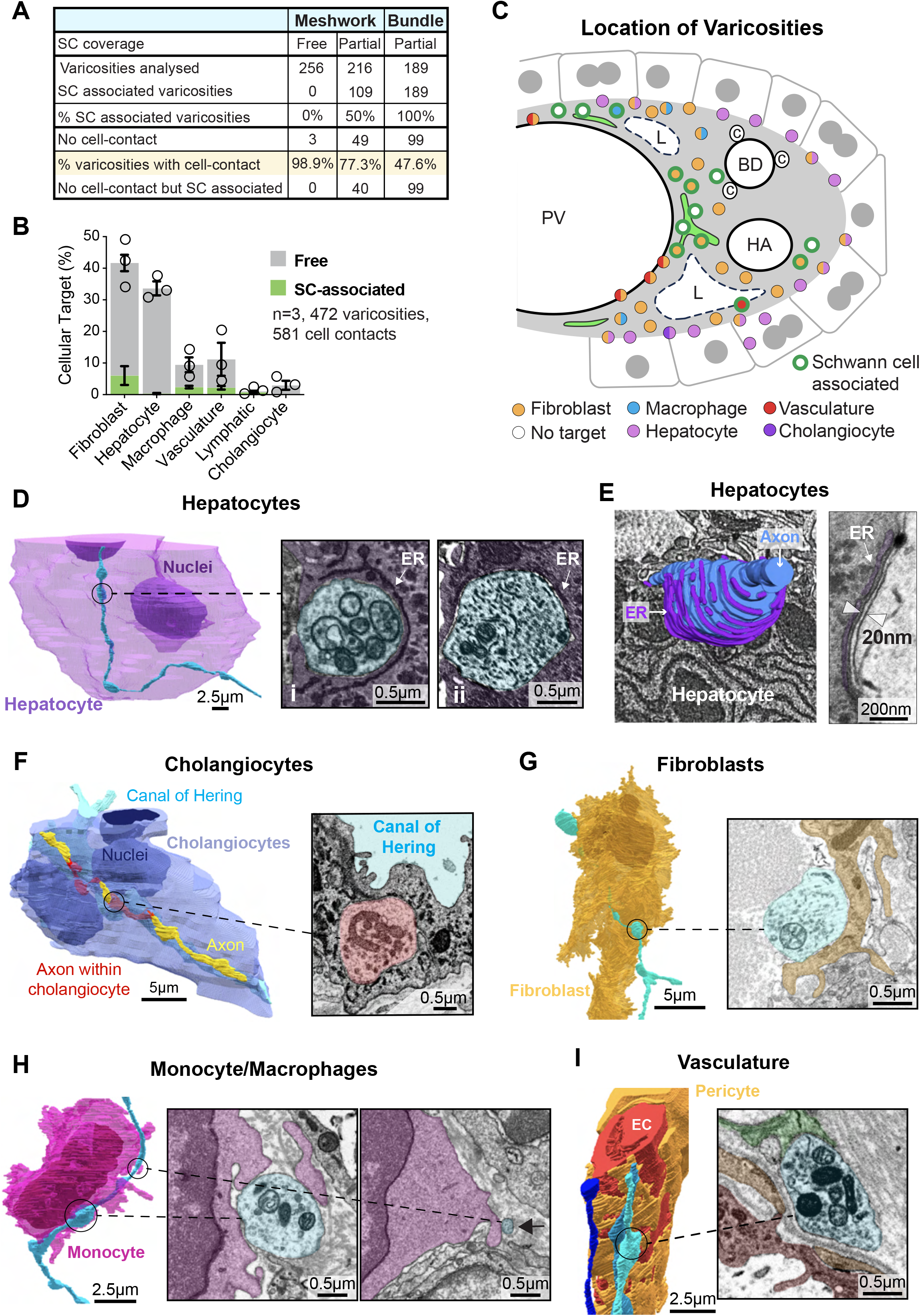
The sympathetic network makes “en passant” connections with multiple cell types. (A) Table shows the proportion of varicosities within the sympathetic network that make contact with a cell within the portal tract. Varicosities are defined as part of the meshwork or within bundles and whether they are associated with SCs (see Table S2 and Methods). (B) Graph shows quantification of the interactions of individual varicosities with a specific cell-type. Varicosities are defined as either SC-associated (green) or free of SC coverage (grey). N=3 mice. Data is shown as mean (circles for each animal) ± SEM for free (grey) and SC-associated (green) bars. (C) Schematic illustrating the spatial distribution of axon varicosities across the portal tract within the analysed datasets. 50 varicosities were randomly selected from Table S2 and their location mapped. (D) Representative images of the interaction of varicosities with hepatocytes. Left image shows a segmented individual axon (blue) traversing a segmented hepatocyte (purple) with the accompanying EM image showing (i) the protrusion of the varicosity (filled with mitochondria and vesicles), into the hepatocyte with alignment of the hepatocyte ER surrounding the varicosity. (ii) varicosity from a second axon. (E) 3D reconstruction of the ER of the hepatocyte (purple) with its associated segmented varicosity (blue) superimposed on its EM image. TEM image shows the “synaptic” space (∼20nm) between a varicosity and the associated hepatocyte. Arrowheads indicate the synaptic space. Arrow indicates aligned endoplasmic reticulum (ER) (see Movie S4). Representative images of the interaction of varicosities with (F) a cholangiocyte (G) a fibroblast, (H) a monocyte/macrophage and (I) the vasculature. Left image shows three segmented axons (blue shades) interacting with a vessel (red) and its pericyte (orange). Associated EM image shows the indicated varicosity associating with the basal lamina surrounding the pericyte of the vessel. Note the varicosity is associated with a SC (green).

Identification of the cell types contacted by varicosities within the meshwork regions revealed that similarly to the terminals, most interactions of the varicosities were not with the vasculature but with fibroblasts throughout the stroma (42%) and hepatocytes at the portal plate (34%) (Figures 6B-C and Table S2). Varicosities were also associated with macrophages, though less frequently (9%). Interestingly, varicosities showed interactions that were not observed for terminals. A small proportion (3%) associated with epithelial cells of both the major bile ducts and with the smaller ductules, known as the Canals of Hering, a proposed niche for cholangiocyte stem/progenitor cells^49^, and a region reported to be regulated by innervation^50^.

Like terminal contacts, varicosity contacts appeared direct and synaptic-like (Figures 6C-I). Also, similarly to the terminals, the nature of the interactions appeared to differ between the cell types. The mostly SC-free hepatocyte contacts frequently involved the protrusion of the varicosity into the hepatocyte, forming an invagination that covered most of the surface of the varicosity (Figure 6D and Movie S4), with a uniform ∼20nm gap between the membrane of the varicosity and the hepatocyte (Figure 6D, S5E). Consistent with synaptic-like signalling at these contact sites, the varicosities were rich in mitochondria and vesicles of varying shapes and densities polarised towards the site of contact. Moreover, the hepatocyte appeared polarised to the contact, with the ER enwrapping the protruding varicosity indicative of a signalling contact (Figure 6E and Movie S4).

Interestingly, the rarer contacts observed with cholangiocytes, a second hepatic epithelial cell type, were mostly less direct involving interactions with the basal lamina of the cholangiocytes (Figure S5E-F). However, further sampling of cholangiocytes at the Canals of Hering detected interactions similar to those involving hepatocytes, with varicosities protruding into the cell and with structures indicative of direct synaptic-like signalling (Figure 6F).

In contrast, fibroblast interactions most frequently involved thin protrusions that partially enwrapped a varicosity. The organelles within the varicosities also appeared polarised to the site of interaction, whereas any organisation within the fibroblast was less obvious (Figure 6G and Movie S4). Rarer examples were found close to the nucleus, creating indentations to the shape of the fibroblast (Figure S5G-H), possibly reflecting a distinct form of signalling. Macrophages also made direct contacts with varicosities, (Figure 6H) but we also identified contacts with the axons between the varicosities (Figure 6H) perhaps indicative of the “scanning” role of macrophages observed in other tissues^51–53^. As for the terminals, the interactions of the internal varicosities with the vasculature were somewhat different, in that the interactions were not direct but mostly aligned along the basal lamina and were more frequently associated with SCs (Figure 6I and Figure S5E). Many varicosities (32%) made direct contact with more than one cell type (Figure S6A). The majority of these bivalent contacts involved fibroblasts and hepatocytes but also interactions of either cell type with macrophages. (Figure S6B-C). Notably these interactions with hepatocytes were distinct: the varicosities did not protrude into the hepatocyte permitting the varicosity to interact with a second cell type (Figures S6C).

Analysis of the varicosities found within the larger bundles showed a different variety of interactions consistent with the location of these bundles mainly along vascular structures. Here, the majority of varicosity interactions were with the vasculature (32%) or with fibroblasts (53%) with a higher variability between samples dependent on the location of the sampled volume (Figure S6D-F). Many of the interactions with the vasculature were associated with flattened varicosities that appeared polarised to pericyte contacts with apparent folds in the pericyte plasma membrane that somewhat resembled structures observed at the neuromuscular junction^44^ (Figure S6F). Together, these findings show that the majority of synaptic-like interactions within the sympathetic network are mediated by varicosities along the length of the axons, rather than the terminals, with direct engagement of multiple cell types across the portal tract.

### Connectomics reveals a complex circuitry integrating multiple cell types within the portal tract

An advantage of vEM is that it permits the tracing of individual axons within a tissue. Previously, innervation of peripheral tissues has focused on how terminals signal to individual cell types or structures. However, our analysis of the connectome of the liver shows that most of the interactions do not occur at terminals but at the multiple varicosities found along the length of individual axons within a network structure. In principle, these multiple points of contact would allow individual axons to co-ordinate interactions within the liver. To determine the connectome of individual axons, we traced axons from multiple datasets and annotated contacts along their length (Table S2). This analysis revealed a variety of cellular interactions along the length of each axon, but remarkably 100% of the axons analysed (n=60) made direct contact with more than one cell type (Figure 7A and Table S2), revealing the complex interactome of the sympathetic network.

**Figure 7:**
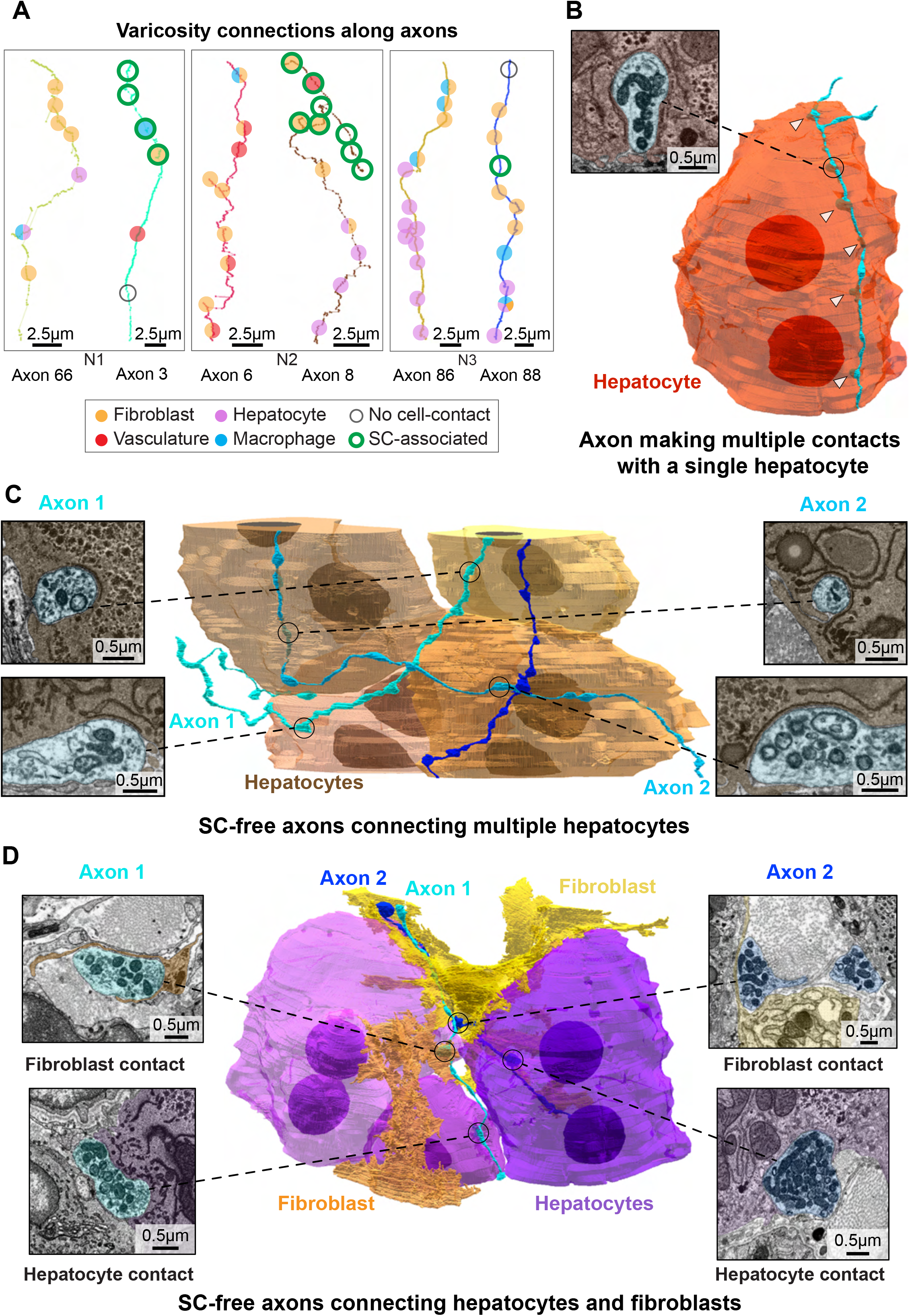
Connectomics reveals local circuitry integrating multiple cell-types within the portal tract. (A) Representative axon skeletons illustrating indicated cell-type specific interactions at sequential varicosities. SC-associated varicosities are indicated by green outline. (B) Image shows an example of a segmented axon (blue) making multiple contacts along a single hepatocyte (red). Varicosities protruding into the hepatocyte are highlighted by arrowheads, with the indicated varicosity shown in the EM image. (C) Image shows segmented axons (shades of blue) connecting neighbouring hepatocytes (shades of orange). Indicated varicosities are shown in EM images. Note the hepatocyte ER is orientated around the varicosities in all examples. (D) Images show segmented axons connecting fibroblasts (yellow) and hepatocytes (purple). Indicated varicosities are shown in EM images.

Interestingly, there were several different patterns of organisation, potentially providing distinct modalities of signalling: (i) Sequential varicosities along an axon making multiple contacts with the same cell; for example, a hepatocyte – that may allow a more robust signal to an individual cell (Figure 7B). (ii) Sequential varicosities contacting neighbouring cells of the same type; for example hepatocytes at the portal plate - potentially co-ordinating signalling between similar cells (Figure 7C) (iii) Sequential varicosities contacting different cell types; this was observed for all axons examined with examples including hepatocytes and fibroblasts (Figure 7D), hepatocytes and cholangiocytes at the Canal of Hering (Figure S7A), and the vasculature and surrounding fibroblasts and macrophages (Figure S7B) - providing a mechanism to co-ordinate different cell types within the liver. Together, this quantitative, connectomic analysis confirmed the integrated network nature of liver innervation, providing a new paradigm of how peripheral tissues are regulated by the nervous system.

## Discussion

The peripheral nervous system (PNS) is essential for regulating tissue function, yet how many peripheral tissues are innervated to control their function remains poorly defined. Here, we have taken a quantitative, cross-scale, 3D-imaging approach to determine the nature of liver innervation. Previous work identified that the liver is primarily innervated by sympathetic neurons associated with vascular structures and bile ducts^18–20^. Our cross-scale, connectomics approach has confirmed the dominancy of sympathetic innervation but has further uncovered a complex neural network that spreads across and along the portal tract, interfacing with multiple cell types within the liver, thereby challenging our perceptions of how peripheral tissues are innervated.

Firstly, we found that while sympathetic contacts within liver do occur at terminals, the vast majority of contacts take place at varicosities along the length of individual axons. The periodicity of these contacts is frequent, similar to the distancing of synapses along axons within the brain^32^, implying an integrated network style of signalling in contrast to the hierarchal, tree-like structures envisioned for the innervation of peripheral tissues. The network structure is further confirmed by the finding of axons running in opposite directions in small bundles and that individual axons contact multiple cell types along their lengths. These findings imply an integrative network, in which signalling is distributed along the length of axons rather than confined to terminal structures, but further work is required to determine how this network controls liver function.

Secondly, we found that most sympathetic contacts were not with the vasculature but instead involve direct contacts with hepatocytes and stromal cells. At the structural level, our analyses showed that the varicosities were full of mitochondria and vesicles directed towards the site of the cellular interactions and that the directly contacted cells appeared to polarise in response to these interactions, with ER structures aligned to the point of contact, consistent with a synaptic-like junction^54^. However, these interactions were quite distinct between the different cell types, with varicosities penetrating into hepatocytes, whereas the fibroblast interactions mostly involved protrusions wrapping around exposed varicosities. These differences suggest that the nature of axon-cell interactions is tailored to the identity of the target cell, potentially reflecting distinct modes of communication within the network.

Sympathetic neurons release nor-adrenaline and it has been reported that hepatocytes and stromal cells express different combinations of the cognate receptors and can respond to these signals^55^. Moreover, sympathetic neurons often co-secrete NPY and express neuropeptides that may act distinctly on the different cell types^56,57^. Recent single cell analyses, chemogenetic and retrograde tracing studies of sympathetic ganglia have revealed distinct sub-sets of neurons, which differentially innervate different tissues, suggesting the existence of tissue-specific guidance trajectories and molecular programmes that mediate interactions with specialised cellular environments^33,58–61^. However, our findings that individual axons contact multiple cell types along their length, may indicate a certain promiscuity in cellular interactions that would theoretically permit the establishment of a flexible network structure to control and co-ordinate tissue function. In addition to afferent signalling, sympathetic neurons respond to neurotrophins such as NT3 and NGF, the latter of which can be trafficked back to the nucleus^33^. This raises the possibility that bi-directional signalling permits the sympathetic network to both sense and control liver function. In future work it will be of great interest to determine how these signalling structures control liver biology within the context of the organism.

Thirdly, we found that in contrast to current dogma, much of the axonal network is devoid of SC coverage. Schwann cells have been reported to envelop all peripheral axons with rare exceptions associated with terminal structures within the skin and cornea^24,40,43^. This coverage has been considered essential for providing metabolic and other nurturing support to the axons of peripheral neurons that can extend for long distances from the neuronal cell body^39^. The liver is innervated by fibres originating from the superior mesenteric and celiac peripheral ganglia, that reside outside of the liver^58,60^. That much of this network is free of SC coverage is remarkable and shows that neurons can cope with extensive stretches of their axons devoid of glial coverage. Moreover, the axonal network is also not covered by other cell types that might provide a protective barrier to the axons, such as perineurial fibroblasts. This lack of any observable blood nerve barrier (BNB)^62^, also indicates these axons are stable without this protection. Interestingly, we observed that SC protrusions appeared to be organised around some varicosities in mostly SC-free axons; this included mitochondria proximity and the alignment of the ER. It is an enticing possibility that these are sites where SCs can still “nurture” axons at distances from the ganglia.

Despite this apparent resilience, recent work has shown that the sympathetic axonal network is highly sensitive to metabolic challenge, resulting in large-scale loss of the sympathetic nerves within the liver^18,19,22^, which may be due to either a lack of glial coverage or existing in the absence of a BNB. Interestingly, the loss of the axonal network is reversible, recovering upon removal of the metabolic stress. Our previous work has shown that axonal regeneration is critically dependent on SCs to guide regrowing axons back to their target^26,42,63,64^, so it is of great interest how this network may regenerate in the absence of associated SC.

In summary, this work describes a quantitative, cross-scale 3D imaging approach to define the connectome of the liver, that should be transferable to the analysis of other peripheral tissues and sites of new innervation, such as during tumour formation^64,65^. Here, it has resulted in the discovery of a complex sympathetic network that connects different cell types across the portal tract of the liver with many parallels to networks in the CNS. Future work will define how this network acts to control liver function in both a healthy organism and its role in metabolic and other diseases.

## Resource Availability

### Lead contact

Further information and requests for resources and reagents should be directed to and will be fulfilled by the lead contact, Alison Lloyd.

### Materials availability

This study did not generate new unique reagents.

### Data and code availability

All data reported in this paper will be shared by the lead contact upon request. VolumeEM datasets are publicly available on Webknossos, links can be found in Supplementary Table 1.

## Acknowledgments

This work was supported by CRUK (C378/A28282); Generalitat Valenciana (CDEIGENT, CDEI-05/20-C); Gobierno de España Ministerio de ciencia y Innovación, (AEI; PID2020-117879GB-I00/AEI/10.13039/501100011033); Swedish Research Council (VR 2023-01847); Swedish Foundation for Strategic Reserach (SSF-RIF2); Swedish National Microscopy Infrastructure (NMI, VR-RIF2023-00163); Ragnar Söderberg Foundation (S11/20); Novonordisk Foundation (NNF25OC0105515); EU Marie Skłodowska-Curie grant No. 101063809 and the CIPF–ISABIAL 3D-Electron Microscopy Unit. The UCL LMCB EM facility (RRID:SCR_027340) is supported by the Wellcome Trust (218278/Z/19/Z) and UKRI via UCL Research Capital Investment Fund 24/25 (CEF7). We thank J.M. García Verdugo and Andrea Ibanez-Grau for advice on EM, Andreas Mueller for animation/Blender support, the annotation team listed in methods, UCL/CIPF Biological Services, and the Lloyd lab for useful discussions.

## Author contributions

Conceptualisation: L.N., J.B. and A.C.L. Development of Methodology: J.B. Investigation: L.N., G.C. and J.B., with help from M.S.N., A.L.C., R.C., C.A.U., C.R.S., A.L.T., J.F.R., E.V., J.A.G.S., M.D., A.P., A.R. and E.R.A. Formal analysis and writing: L.N., G.C., J.B. and A.C.L. Supervision: A.C.L.

---

**Figure S1, related to Figure 1:**
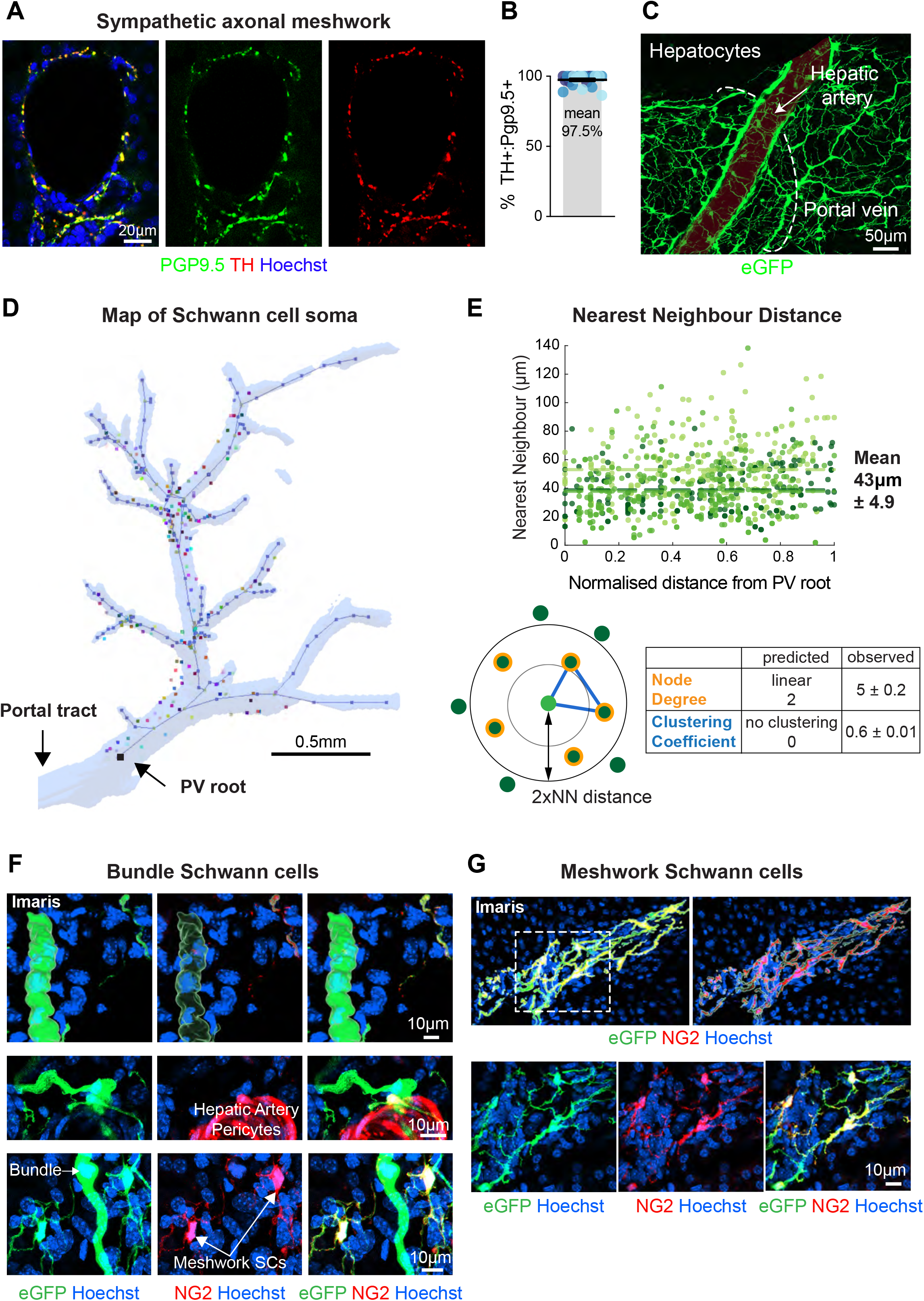
The portal tract is innervated by a meshwork of sympathetic neurons. (A)Representative confocal images of the portal tract of mouse liver immunolabelled to detect axons (PGP9.5, green), sympathetic axons (TH, red), and nuclei (Hoechst, blue). (B) Quantification of (A) showing the percentage of axons that are sympathetic (n=3 mice, shades of blue). Data shown as mean ± SEM. (C) Representative confocal image of the eGFP meshwork within the portal tract. (D) 3D segmented reconstruction of light-sheet image (Figure 1B), converted into a vectorised representation of the portal tract, with the proximal reference point defined as the portal vein (PV) root and SC soma. (E) Nearest-neighbour analysis of SC soma distribution plotted as a function of distance from the PV root. The mean nearest-neighbour distance (43µm) remains relatively constant along the length of the portal tract (n=3, shades of green). 3D analysis of clustering coefficients indicates a 3D meshwork structure rather than a linear organisation. (F) Representative confocal images immunolabelled to detect NG2 (red), SCs (eGFP, green) and nuclei (Hoechst, blue). Images show that SCs in nerve bundles do not express NG2. Top image shows Imaris surface rendering of the bundles. Middle image shows NG2+ pericytes and NG2-SCs (green) in the nerve bundles associated with the hepatic artery. Lower images show meshwork SCs that co-label for eGFP and NG2. (G) Representative confocal images of the meshwork region of the sympathetic network immunolabelled to detect NG2 (red), SCs (eGFP, green) and nuclei (Hoechst, blue). Top image shows Imaris definition of the surface of the SCs and show co-labelling of NG2. Lower image shows the distinct ramified appearance of meshwork SCs.

**Figure S2, related to Figure 2:**
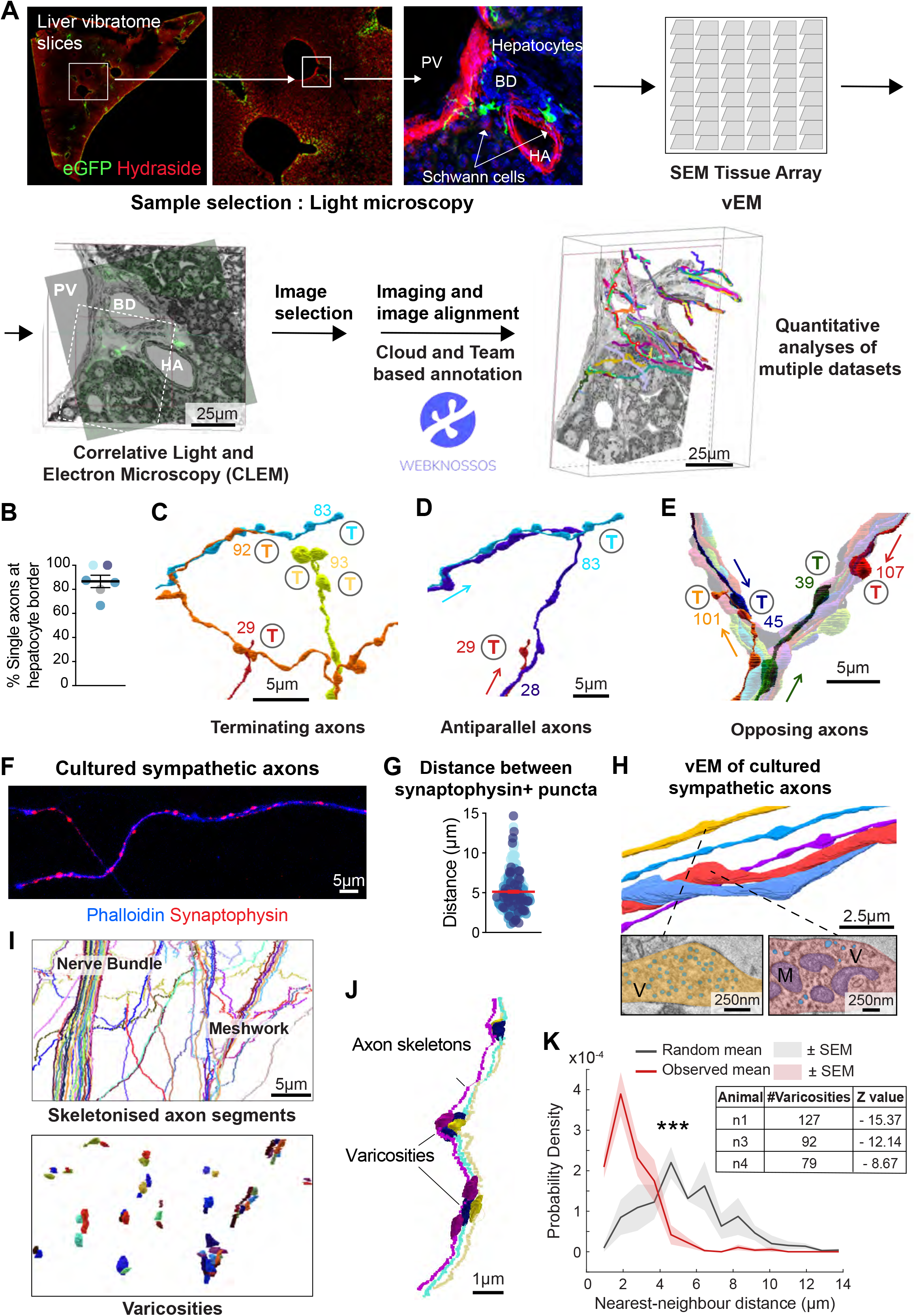
Volume EM reveals a complex axonal network within the portal tract. (A) Workflow for quantitative, vEM to analyse the innervation of mouse liver. Fluorescence of eGFP expressed by SCs was used to identify regions of interest within vibratome sections of mouse liver from *PLP-eGFP* mice (portal vein (PV), bile duct (BD), and hepatic artery (HA). Excised regions were subsequently sectioned for Array Tomography (AT). CLEM was used to direct imaging to regions of interest, and the resultant 3D datasets acquired by scanning electron microscopy (SEM) were aligned and annotated using cloud and team-based annotation workflows. Quantitative methods were developed to analyse the multiple datasets in a non-biased fashion (see Movie S2, Methods for details and Tables S1 for complete datasets). (B) Graph shows % of single axons at the hepatocyte border. N=4, shades of blue, mean ± SEM. (C) Segmented reconstruction of the axons shown in Figure 2B that terminate within this sub-group. T=terminal. (D) Example from Figure 2B illustrating axons running in opposing directions within the same bundle. Axon 28 (purple) traverses between two axon bundles containing axons that terminate in opposing directions (axons 29 (red) and 83 (blue)), indicating that axons can run antiparallel within the same bundle. (E) Example from the broader dataset showing axon terminals converging within bundles, illustrating the bidirectional organisation of axons within the portal tract network. (F) Representative confocal image of cultured mouse sympathetic axons immunolabelled to detect actin (phalloidin, blue) and synaptophysin (red). (G) Graph shows the distance between synaptophysin puncta within individual axons (n=3 explants). Red line shows mean ± SEM. (H) Example of a vEM reconstruction of cultured sympathetic axons. Dashed line indicates EM images, highlighting the accumulation of mitochondria (M) and vesicles (V) within varicosities. (I) Example of skeletonised axons within the sympathetic network (top image) and their segmented varicosities (lower image), highlighting clustering of varicosities in the meshwork. (J) Example of skeletonised axons and their varicosities, highlighting clustering of varicosities within a bundle. (K) Graph shows quantification of varicosity clustering analysis as assessed by comparing the distribution of nearest-neighbour distances (NNDs) between varicosities across three independent datasets (red, mean ± SEM; n=3 mice), compared to simulated random point distributions generated within the same spatial domain (grey, mean ± SEM; 999 simulations per dataset). The observed distribution indicates clustering of varicosities. Shaded areas represent SEM across datasets. Table shows the number of detected varicosities per dataset and corresponding z-scores comparing observed mean NNDs with simulated null distributions. ***p=0.001 by one-sided Monte Carlo randomisation test.

**Figure S3, related to Figure 4:**
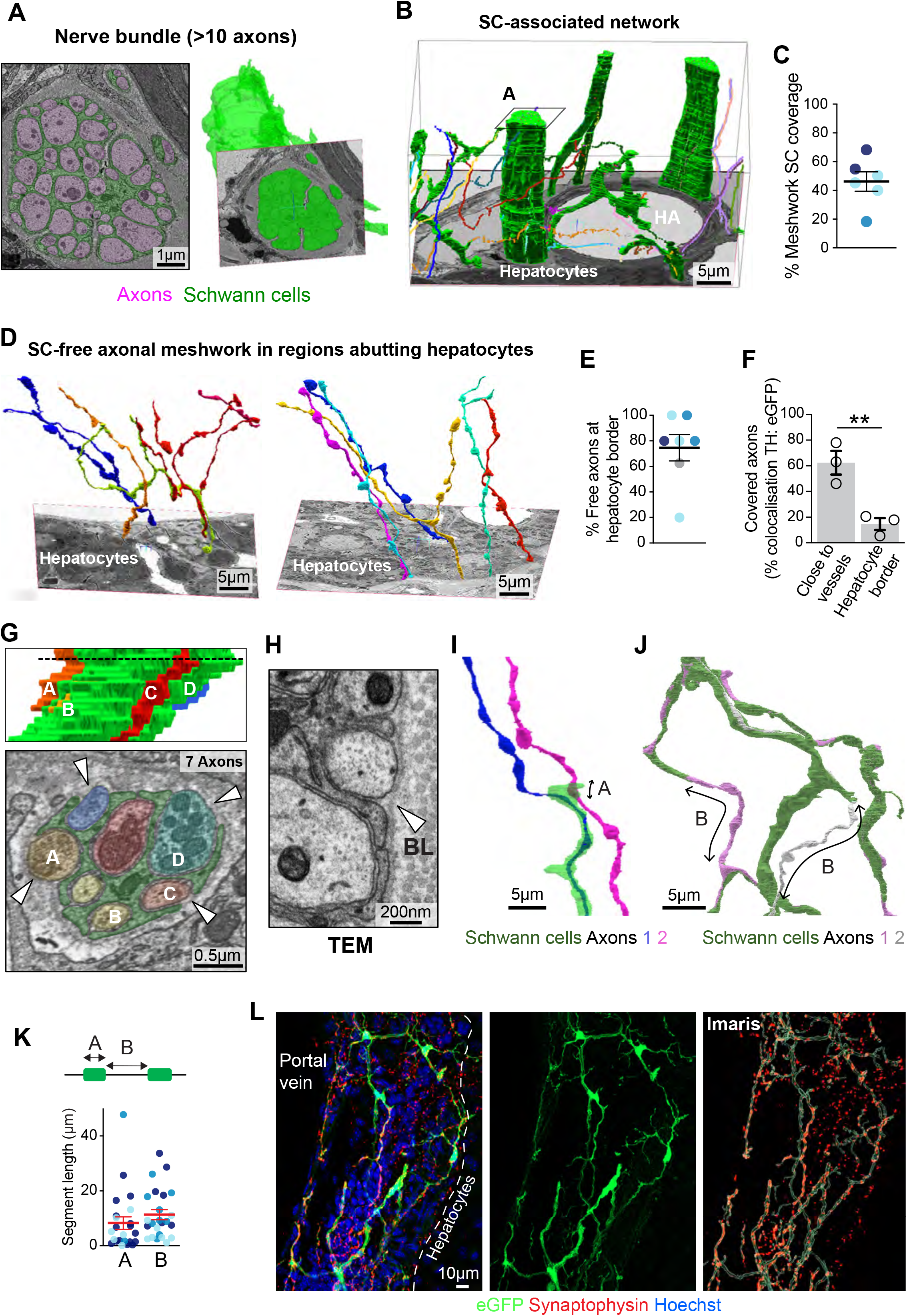
The axonal network of the portal tract is only partially covered by Schwann cells. (A) The EM image (left) shows a classical Remak bundle organisation of a large bundle, where axons (pink) are surrounded by a SC (green). The reconstructed volume (right) shows the bundle (green). (B) 3D volume containing the bundle in (A) in which the SCs have been segmented to show coverage of large and small bundles as well as SC-free axons within the reconstructed region. (C) Graph shows % SC coverage of the meshwork, excluding bundles. N=3, shades of blue, mean ± SEM. (D) EM volumes showing segmented free single axons that lack SC association, forming a meshwork at the hepatocyte border shown in superimposed EM sections. (E) Graph shows % of free axons at the hepatocyte border. N=4, shades of blue, mean ± SEM. (F) Quantification of Figure 4C showing the % of SC coverage of sympathetic axons in different portal tract regions. N=3 mice, mean ± SEM. (G) Example of a bundle of seven axons exhibiting partial SC coverage while remaining enclosed within the SC basal lamina. The segmented reconstruction (top) highlights axons A-D with partial SC coverage. EM slice (bottom) with arrowheads indicating exposed regions that remain within the SC basal lamina. (H) TEM zoom of a partially exposed bundle showing axons remain enclosed with the basal lamina (BL). (I) 3D image of 2 segmented axons in which a SC (green) protrusion makes brief contact with an exposed axon. (J) 3D reconstruction showing regions of exposed axon between sequential SC contacts. (K) Graph shows (A) Length of SC contact (see G) and (B) Length between sequential SC contacts (see H). N=3, shades of blue, red lines show mean ± SEM. (L) Representative confocal and Imaris reconstructed images of the portal tract immunostained to detect synaptophysin (red), SCs (eGFP, green) and nuclei (Hoechst, blue). Note synaptophysin staining in regions free of SC coverage. **p<0.01 by unpaired t-test.

**Figure S4, related to Figure 5:**
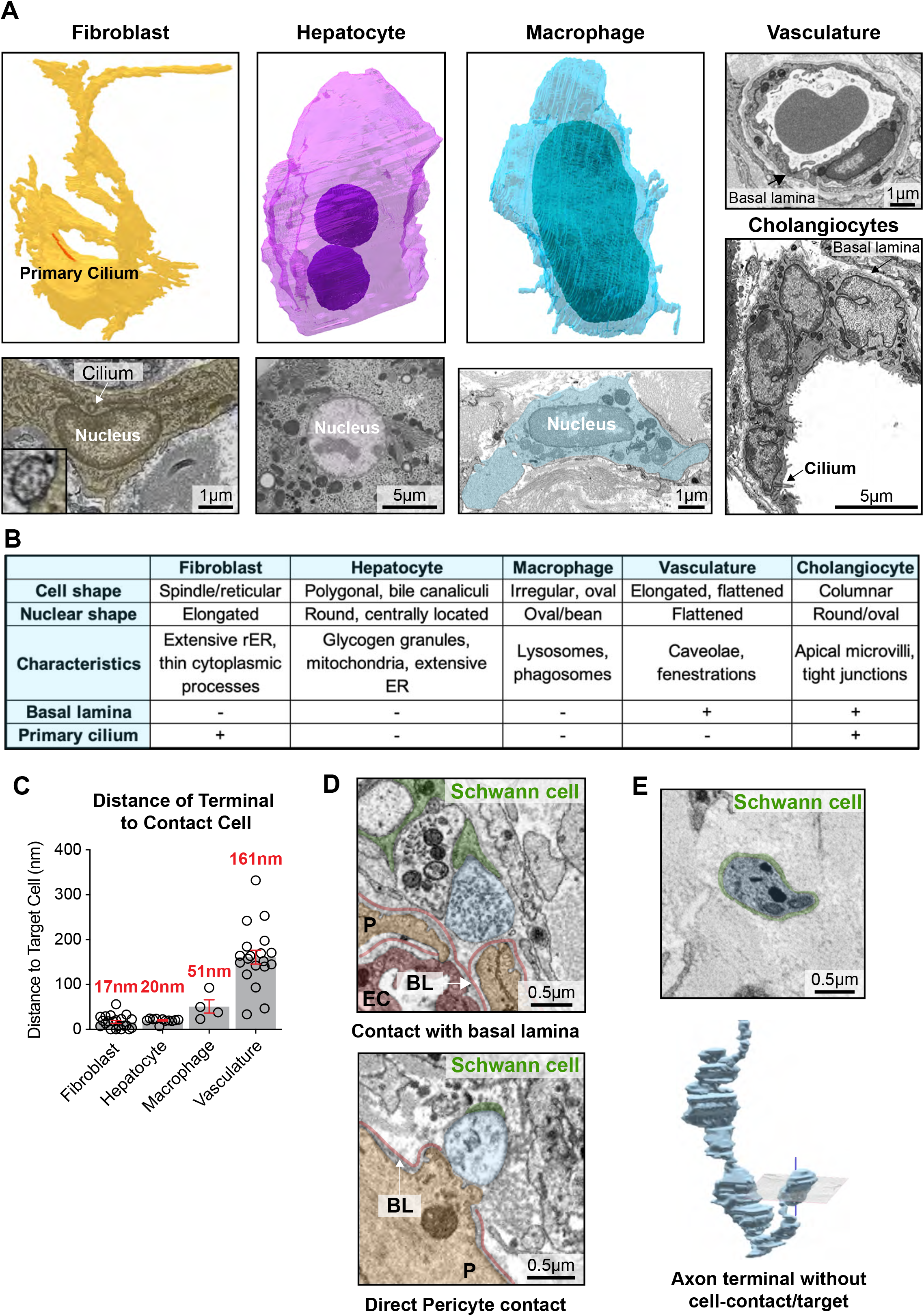
The sympathetic network terminals contact distinct cell types within the liver: (A) Representative images of the indicated cell types that permit their identification within EM images. See Methods for full details. (B) Table defining the ultrastructural characteristics of each cell type found within the portal tract. (C) Graph shows the quantification of the distance between terminals and the contacting cell membrane. Each circle represents one terminal. Data shown as mean (red) ± SEM. (D) EM images showing terminals contacting the vasculature at either the basal lamina or making direct contact with a pericyte and are pseudo-coloured to show SCs (green), axons (blue), pericytes (P, orange), endothelium (EC, red) and basal lamina (BL, red). (E) EM image of a terminal that does not make contact with a cell. Lower image shows a 3D rendering of the terminal.

**Figure S5, related to Figure 6:**
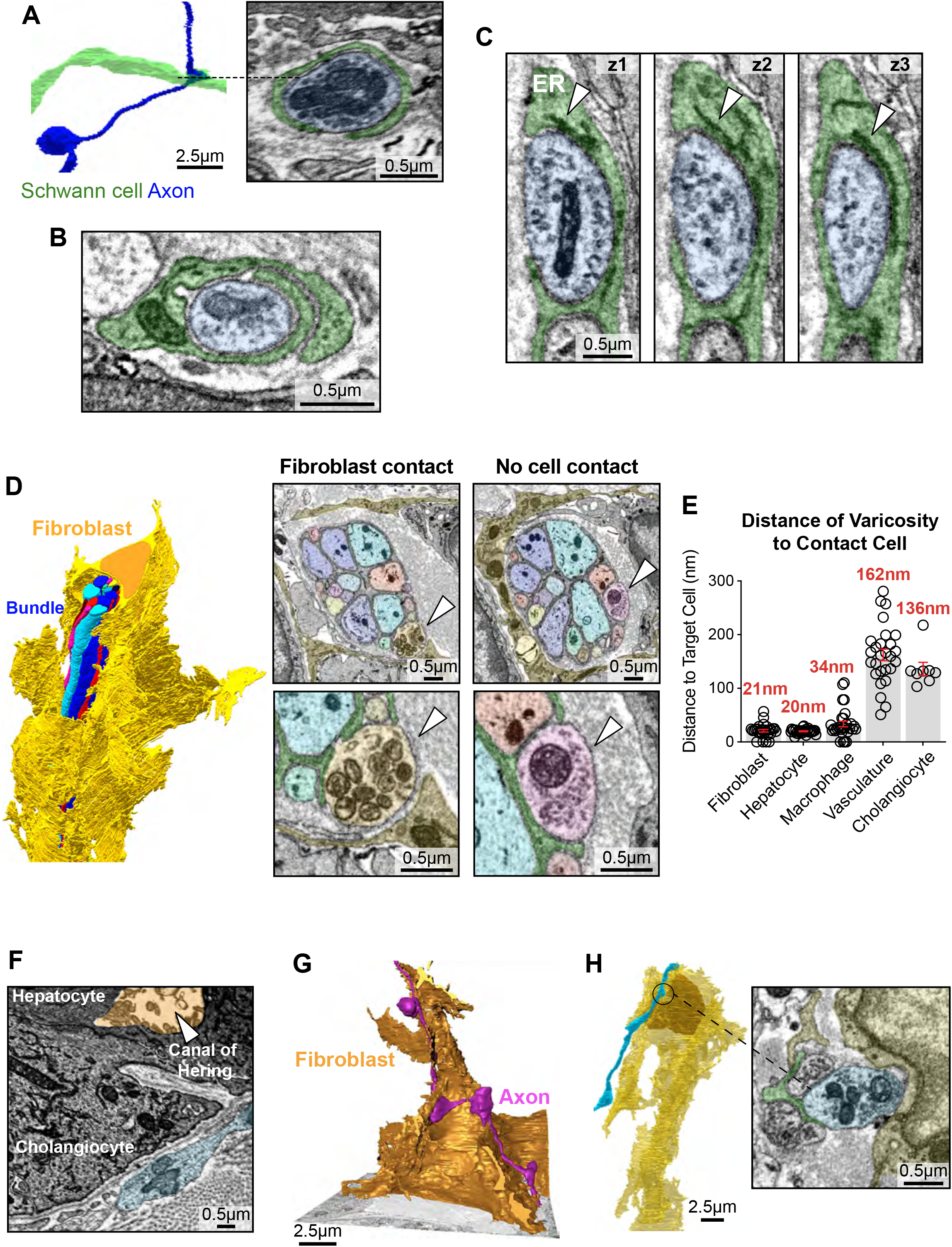
The sympathetic network makes “en passant” connections with multiple cell types. (A) Segmentation of an axon and an associated SC, the indicated EM image showing a varicosity (blue) enwrapped by a SC (green) in the absence of a separate cell interaction. (B) EM image of a separate example of a SC (green) surrounding a varicosity (blue). (C) EM image series (z1-3) showing a varicosity enwrapped by a SC (green) with the ER (arrowhead) aligned to the varicosity. (D) Left image shows a segmented bundle surrounded by a fibroblast (yellow). Right images show EM images of cross sections along the length of the bundle showing varicosities (arrowheads) contacting a fibroblast (yellow) or no cell but instead exposed to the extracellular matrix. (E) Graph shows the quantification of the distance between varicosities and the contacting cell membrane. Each circle represents one varicosity. Data shown as mean (red) ± SEM. (F) EM image of the interaction of a varicosity (blue) with a cholangiocyte at the Canal of Hering (orange). (G) Segmented axon (pink) with prominent varicosities, interacting with a fibroblast (yellow) (see Movie S4). (H) Left image shows a segmented axon (blue) making contact with a segmented fibroblast (yellow). EM image shows the indicated varicosity contacting a fibroblast close to the nucleus. Note the varicosity is associated with a SC (green).

**Figure S6, related to Figure 6:**
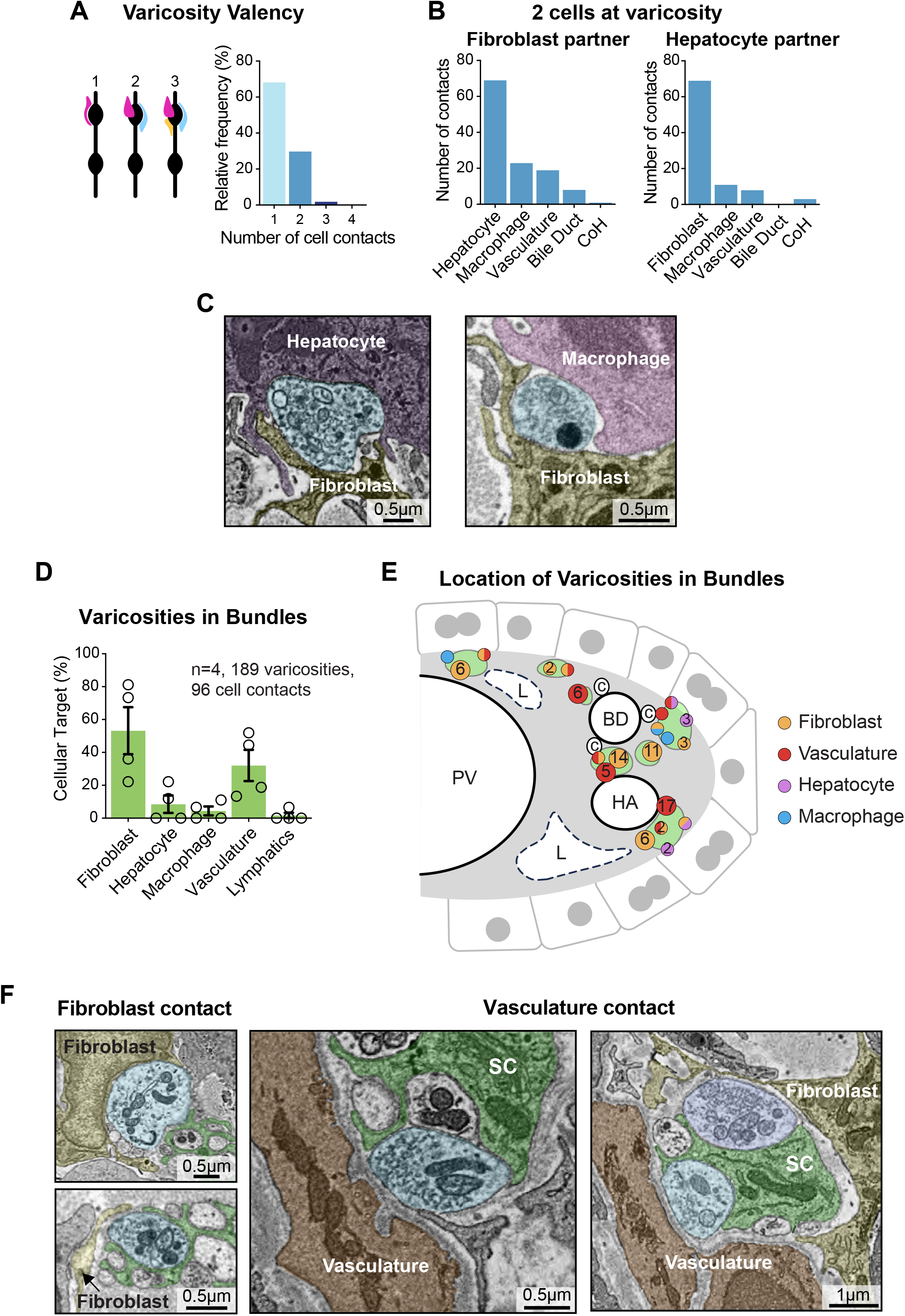
The sympathetic network makes “en passant” connections with multiple cell types. (A) Schematic representation and quantification of the number of cell contacts at individual varicosities (484 varicosities from n=4 mice). (B) Quantification of the relative frequency of partners at varicosities with fibroblasts (left, 120 varicosities) and hepatocytes (right, 91 varicosities). (C) Representative EM images of varicosities contacting two cell types, pseudo-coloured to show axons (blue) and fibroblasts (yellow), hepatocytes (purple) and macrophages (pink). (D) Graph shows quantification of the interactions of varicosities within bundles with specific cell types. N=4 mice, mean ± SEM. (E) Schematic illustrating the spatial distribution of the cell-contacts associated with axon varicosities in bundles across the portal tract. (F) Representative EM images of varicosities within bundles contacting a fibroblast (yellow) close to its nucleus or at a protrusion and contacting the vasculature, with varicosities making contact with the basal lamina at the surface of pericytes (brown).

**Figure S7, related to Figure 7:**
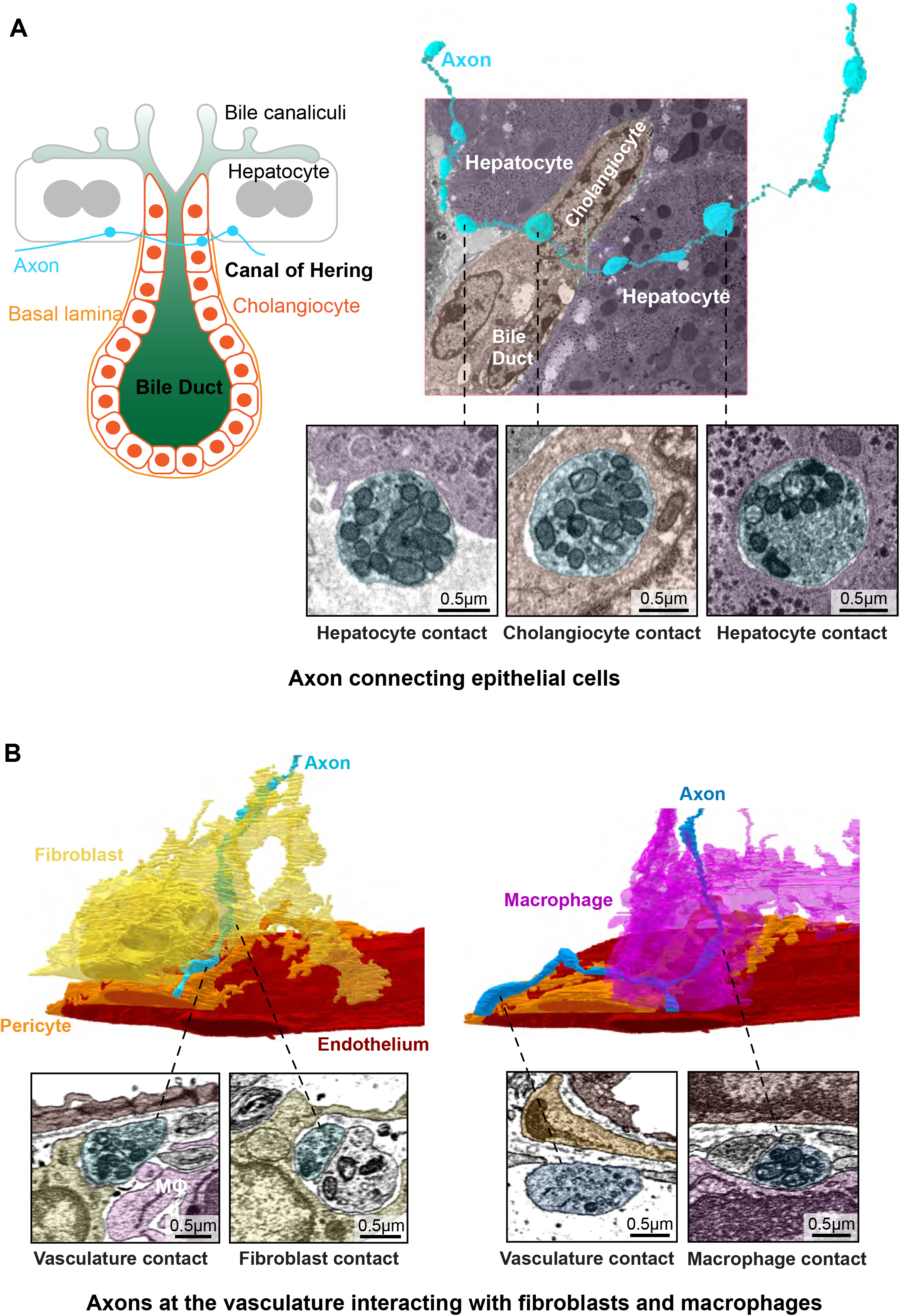
Connectomics reveals local circuitry integrating multiple cell types within the portal tract. (A) Schematic illustrating an example of the contacts of an axon at the bile duct. Image shows a segmented axon (blue) superimposed on an EM image, making contacts with hepatocytes and a cholangiocyte surrounding the bile duct. Indicated varicosities show zoomed EM images, sequentially contacting a hepatocyte, a cholangiocyte and a second hepatocyte. (B) Images show segmented axons with sequential varicosities interacting with the vasculature (endothelial cells, red and pericytes, orange), and a fibroblast (yellow, left) and macrophage (pink, right). Indicated varicosities are highlighted in EM images. Note the varicosities contact the vasculature at the basal lamina.

## Supplemental Table Legends

**Table S1. Summary of vEM datasets used in this study.**

Summary of all vEM datasets generated in this study, including animal number, dataset reference, number of serial sections, reconstructed ROI dimensions, tissue volume (calculated by subtracting the estimated PV volume), total annotated axon length, annotation status, axon density and description of cell types within the region.

**Table S2. Axonal Varicosities and Terminals.**

Dataset of axon varicosities and terminals showing; animal and dataset identifiers, axon reference, spatial coordinates, SC association, axon length, and associated target cell types. Each row represents a single annotated terminal or varicosity. Cell-contact annotations are coded according to SC association, where 1 represents non-SC-associated and 2 represents SC-associated. T* represents blunt terminals, whereas T represents bulbous terminals.

## Supplemental Movie Legends

**Movie S1. 3D architecture of liver sympathetic innervation throughout the portal tract, related to Figure 1B**: Reconstruction of light-sheet images of an iDISCO-cleared Liver lobule from *PLP-eGFP* mice immunolabelled to detect Tyrosine Hydroxylase (red) and expressing eGFP (green).

**Movie S2. Volume EM of the liver portal tract, related to Figure S2A:** 3D EM reconstruction of a representative region of the portal tract showing the axonal meshwork and SC coverage. Movie show skeletonised axons together with segmented SCs (green) within the volume.

**Movie S3. Free meshwork at the hepatocyte border, related to Figure 4B**: 3D EM reconstruction of a representative example of the portal tract at the hepatocyte border showing SC-free axons.

**Movie S4. Varicosities associate with hepatocytes and fibroblasts, related to Figure 6E and S5G:** 3D EM reconstruction of a representative example of a varicosity (blue) associated with a hepatocyte, with the hepatocyte segmented in pink with its ER shown in purple. The second example shows a representative varicosity (pink) associated with a fibroblast (orange).

**Movie S5. Individual axons contact multiple cell types, related to Figure 7D**: Connectomic analysis shows representative examples of individual axons contacting multiple cell types. The first example shows axon 99 sequentially associating with a hepatocyte, a fibroblast and a second hepatocyte. The second example shows axon 88 sequentially contacting a hepatocyte, macrophage, and a fibroblast.

## METHODS

### Experimental Model Details

#### Animals

All animal work was performed in accordance with UK Home Office guidelines at the Central Biological Services Unit at University College London (UCL, UK), or in accordance with institutional and regional ethical approvals at the Centro de Investigación Príncipe Felipe (Valencia, Spain) and authorised by the Generalitat Valenciana. The following transgenic mouse reporter lines were used in this study: *Plp-eGFP* mice in which all Schwann cells (SC) express eGFP^25^, and *Ng2-*reporter mice, in which a red fluorescent protein variant (DsRed.T1) is expressed in NG2-expressing cells^29^. Mice and rats were housed in pathogen-free individually ventilated cages under controlled temperature (20-24°C) and humidity (50-60%) on a 12h light/dark cycle with ad libitum access to food and water. Both male and female adult mice (8-24 weeks of age) were used.

### Method Details

#### Neuronal sympathetic cultures

Superior cervical ganglia (SCG) were dissected from post-natal day 1 (P1) Sprague Dawley rats of both sexes and were enzymatically dissociated and plated onto 13mm glass coverslips (30000cells/coverslip) coated with collagen and laminin (1:200) ^66^. Cultures were maintained for 6 days in DMEM containing 10% Fetal Bovine Serum, 2mM glutamine, 1% penicillin / streptomycin antibiotics, 10μΜ AraC and 100ng/ ml NGF, at 37°C with 10% CO_2_. 10μΜ AraC was re-added to the medium on the second day until the end of the experiment. The culture medium was refreshed every 2-3 days. SCG explants were washed with PBS and fixed using 4% PFA/PBS at RT for 10mins.

#### Tissue harvesting and fixation

All mice were terminally perfused following deep anaesthesia induced by intraperitoneal injection of pentobarbital (500mg/kg). Where indicated, mice received 1mg/ml Rhodamine-conjugated *Griffonia Simplicifolia* lectin (Vector Laboratories) via the left ventricle for 2 minutes prior to perfusion-fixation to label the vasculature. Transcardial perfusion was performed via the left ventricle using ice-cold fixative delivered through a 25-gauge Luer-lock butterfly needle (Venofix) connected to a peristaltic pump (Gilson Minipuls 3) at a flow rate of 4 mL/min for ∼10 min. Fixative composition was selected according to the downstream application: 4% paraformaldehyde (TAAB Laboratories) in phosphate-buffered saline (PBS; pH 7.4, without Ca²⁺ and Mg²⁺) for fluorescence-based imaging, while a mixture of 2% PFA/2% glutaraldehyde (Electron Microscopy Sciences) in PBS was used for electron microscopy and correlative light and electron microscopy (CLEM) to optimise ultrastructural preservation. Following perfusion, the left liver lobe or the entire liver was harvested and post-fixed overnight at 4°C in the corresponding fixative. Samples were washed in PBS prior to downstream processing.

#### Tissue clearing

For whole organ imaging, liver samples were processed using an iDISCO+ protocol adapted from^67^. Briefly, 4% PFA-fixed liver was dehydrated through a graded methanol series (20-100%) and bleached overnight in 5% hydrogen peroxide in methanol at 4°C. Samples were then rehydrated for whole-mount immunolabelling (see Immunofluorescence section), followed by dehydration in methanol prior to clearing for 3 hours in 66% dichloromethane (DCM) / 33% methanol, and two 30 min washes in 100% DCM for lipid removal. Cleared tissues were transferred to dibenzyl ether (DBE) for refractive index matching and storage prior to light-sheet microscopy.

For confocal imaging of thicker sections (200µm) (Figure 1C), passive clarity technique (PACT) clearing method was performed^68^. Briefly, the PFA-fixed liver tissue was vibratome sectioned into slices (Leica VT1200 S) and incubated in hydrogel solution containing 4% (w/v) acrylamide and 0.25% (w/v) 2,2′-azobis(2-methylpropionamidine) dihydrochloride (Sigma-Aldrich) in PBS. Samples were degassed for 5 minutes by nitrogen sparging and polymerised at 37°C for 8 hours with gentle rotation. Sections were cleared in 8% (w/v) SDS in PBS at 37°C for 4 days, with gentle rotation, then incubated in 13.3% (w/v) Histodenz (Sigma-Aldrich) in PBS for 24 hours before imaging of endogenous eGFP and labelled vasculature (Rhodamine).

#### Immunofluorescence

For cryosection-based immunofluorescence, fixed liver tissue was cryoprotected in 30% sucrose in PBS overnight at 4°C, embedded in OCT, and sectioned (40µm) using a cryostat (Leica). Sections were permeabilised and blocked in 0.3% Triton X-100 in PBS containing 10% normal goat serum (NGS) for 2 hours at room temperature, followed by incubation with primary antibodies overnight at 4°C in PBS containing 10% NGS.

Sections were washed and incubated with Alexa Fluor-conjugated secondary antibodies (1:400) and Hoechst (1:1000) for 1 hour at room temperature. Sections were washed 3 x 15 min in PBS, rinsed in distilled water, and mounted using Fluoromount-G (Invitrogen).

For immunolabelling of cleared tissues, samples processed using the iDISCO+ protocol were incubated in permeabilisation solution (0.2% Triton X-100, 10% DMSO, 6% normal donkey serum in PBS) for 3 days at 37°C following rehydration. Samples were then incubated with primary antibodies against tyrosine hydroxylase (TH; Merck, AB152, 1:150) and GFP (Abcam, Ab13970, 1:300) for 6 days at 37°C in 0.2% Tween-20, heparin (10mg/mL), 5% DMSO, and 3% normal donkey serum in PBS. After overnight washing in PBS, tissues were incubated in secondary antibodies (1:500) for 6 days at 37°C in 0.2% Tween-20, heparin (10 mg/mL), and 3% donkey serum in PBS.

For SCG neurons, the PFA was washed 3x with PBS and the cells were permeabilised and blocked with 0.1% TritonX 100 and 2% normal goat serum in PBS (blocking buffer) for 30min-1h. The primary antibodies were diluted in blocking buffer and incubated overnight at 4°C. The primary antibodies were washed 3x5min with blocking buffer and the cells were incubated with secondary antibodies (1:400) and Phalloidin-546 (Invitrogen A22283, 1:1000) and Hoechst (1:1000), diluted in blocking buffer, for 1h at RT. The secondaries were washed 3x5min with PBS. Coverslips were mounted to slides with Fluoromount-G (Invitrogen).

#### Immunofluorescence antibodies

Antibodies used for immunofluorescence at indicated dilutions: GFP (Chicken, Abcam ab13970, 1:300), PGP9.5 (rabbit, Abcam ab108986, 1:100), Tyrosine Hydroxylase (rabbit, Novus, NB300-109, 1:150), Tyrosine Hydroxylase (sheep, Novus, NB300-110, 1:100), Synaptophysin (Rabbit, Abcam ab32127, 1:100), NG2-PE (rabbit, Abcam, ab314252, 1:1000).

#### Confocal microscopy

Confocal imaging was performed using either a Zeiss LSM900 or Leica TCS SP8 STED with 40x (1.3 NA) and 63x (1.4 NA) oil immersion objectives or a 20x (0.8 NA) objective. PACT-cleared vibratome sections were imaged on a Leica TCS SP8 STED microscope using a 25x (0.95 NA) objective. Z-stacks were acquired with a step size of 0.2 or 1µm, adjusted according to objective magnification and sampling requirements.

For vCLEM workflows, confocal imaging (using both systems described above) was used to identify SC-associated regions of interest (ROIs) based on the endogenous eGFP signal in livers isolated from PLP-eGFP mice.

#### Light-sheet microscopy

Cleared liver caudate lobe samples processed using the iDISCO+ protocol were imaged using a light-sheet fluorescence microscope (Ultramicroscope II, LaVision Biotec) controlled by ImSpector software (LaVision Biotec). Imaging was performed using a 2x objective lens (XLPLN10XSVMP, Olympus; NA 0.6) with an adjustable zoom range (0.63x-6.3x) and an sCMOS camera (Hamamatsu ORCA-Flash4.0). Fluorescence excitation was achieved using 561 nm and 647 nm laser lines. Image stacks were acquired as 16-bit uncompressed TIFF files.

#### Light Microscopy Image analysis

Confocal and light-sheet image datasets were processed and analysed using ImageJ/Fiji ^69^ and Imaris software (Oxford Instruments). For confocal datasets, z-stacks were visualised, cropped and adjusted using Fiji/ImageJ. Maximum intensity projections were generated where appropriate.

Light-sheet datasets were converted into IMS format using Imaris File Converter (Bitplane) and visualised as 3D volumes in Imaris. Rendering and exploration of large volumetric datasets were performed using Imaris 10.0.0 (Bitplane), including generation of optical sections and volume projections. Webknossos software (Scalable Minds) was also used for visualisation and 3D annotation of light-sheet datasets.

For all imaging modalities, adjustments to brightness and contrast were applied across all images within an individual experiment. Final figures were assembled using Adobe Illustrator Software (Adobe Systems). Quantitative analysis of specific features (co-localisation, spatial distribution and morphometric measurements) was performed as described in the relevant sections below.

#### eGFP-Ng2 co-expression analysis

For Figure 1F, quantification of eGFP (Plp-eGFP mice) and NG2 (NG2 immunofluorescence staining) co-expression was performed using FIJI/ImageJ^69^ on confocal z-stack images. SCs were identified based on eGFP expression. Individual SCs were categorised according to morphology into either meshwork or bundle structures. Cell counts were performed manually. The total number of eGFP-positive SC nuclei was determined for each category, and the number of cells co-expressing NG2 was recorded. The percentage of eGFP⁺/Ng2⁺ cells was calculated relative to the total number of eGFP-positive SCs within each category. Analysis was performed on samples from n=3 animals, with 4 independent confocal images analysed per animal and a total of 86 meshwork and 36 bundle nuclei counted.

#### PGP9.5-TH coexpression analysis

For Figure S1A, colocalisation analysis of tyrosine hydroxylase (TH)+ axons with the pan-neuronal marker PGP9.5 was performed on single confocal optical sections using Fiji/ImageJ ^69^. Images were separated into individual fluorescence channels and thresholded independently using fixed intensity thresholds that were maintained across all samples and imaging sessions. Thresholded images were then converted into binary masks. PGP9.5+ axons were identified using the “Analyze Particles” function in Fiji/ImageJ. For each animal (n=3), a total of 236, 248, and 222 individual PGP9.5+ axons were analysed, respectively. TH+ was assessed within each PGP9.5+ region using a fixed mean pixel intensity cutoff that was applied uniformly across all images. The proportion of double-positive axons was calculated as the percentage of PGP9.5+ axons that were also classified as TH+. A minimum of 7 images per animal were analysed from 3 independent animals.

#### Spatial analysis of SCs within the portal tract

For Figure S1D-E, nearest-neighbour (NN) and network analyses were performed on three independent whole-organ light-sheet microscopy datasets obtained from separate animals. Light-sheet fluorescence data were imported into Webknossos^31^ for 3D annotation. SC soma were manually identified based on Plp-directed eGFP fluorescence and annotated using the skeleton tracing tool. In parallel, the branching architecture of the portal vein (PV) was reconstructed using skeleton annotations to provide a positional reference framework for SC soma. A total of 283, 166 and 310 SC soma were analysed across PV skeletons spanning 13249, 4145 and 5776 µm, respectively.

Coordinates of SC soma and PV skeleton nodes were exported from Webknossos as .nml files. Node coordinates were converted to physical units using the voxel calibration stored within the NML files. Pairwise Euclidean distances between all SC soma were calculated in MATLAB R2022b (MathWorks) using “pdist2”. For each soma, the minimum non-self-distance was defined as the NN distance. To determine the position of each SC soma along the portal tract, the PV skeleton was represented as a graph in which annotated skeleton nodes were connected according to the traced PV architecture and weighted by the Euclidean distance between connected nodes. A proximal PV root node, located near the liver hilus at the point of maximal PV diameter within the dataset, was defined as the reference point. The shortest-path distance from the root node to all PV skeleton nodes was calculated, and each SC soma was assigned a position along the PV according to its nearest PV skeleton node. To enable comparison between datasets of different lengths, distances from the PV root were normalised by dividing the shortest-path distance of each soma from the PV root by the maximum PV-root distance within the corresponding dataset, yielding a dimensionless measure of relative position along the portal tract ranging from 0 (PV root) to 1 (most distal soma).

To examine the spatial organisation of SC soma, neighbouring cells were identified in three dimensions. Two SC soma were considered neighbours if they were located within a distance equal to 2x the mean NN distance for the corresponding dataset. Using this criterion, connections were assigned between neighbouring SC soma to generate a network describing local cellular organisation^70^. For each SC soma, the number of neighbouring cells (node degree) was calculated. For example, a node degree of 2 would be expected for a simple linear chain of cells, whereas higher node degree values indicate increased local connectivity within a mesh-like network. In addition, the clustering coefficient was calculated for each SC soma as a measure of how interconnected its neighbouring cells were. A clustering coefficient of 0 indicates that neighbouring cells are not connected to one another, whereas a value of 1 indicates that every neighbouring cell is connected. Node degree and clustering coefficient were calculated for each dataset and represented as mean ± SEM, to quantify the degree of local network organisation.

#### Synaptophysin puncta distance analysis in cultured sympathetic axons

For Figures S2F-G, immunofluorescence images of cultured rat sympathetic axons labelled with phalloidin and synaptophysin were acquired at identical magnification and imaging settings across all samples. Three independent batches (n=3) were analysed, with 3-4 fields of view acquired from 2-3 explants from each batch. Images were analysed in Fiji/ImageJ. For each image, individual axons containing synaptophysin-positive puncta were identified, and the distance between adjacent synaptophysin puncta along the axon was measured manually using the Fiji line tool. Between 6 and 14 inter-puncta distances were recorded per image, a total of 210 distances across the 3 explants was measured. Measurements were performed on all clearly resolvable puncta within the selected field of view.

#### Schwann cell coverage analysis

For Figures 4C and S3F, the association of eGFP-labelled Schwann cells (SCs) with tyrosine hydroxylase (TH)-positive sympathetic axons was quantified from confocal z-stack images using Fiji/ImageJ^69^. ROIs were manually defined based on Hoechst nuclear staining and regions adjacent to blood vessels and regions located at the hepatocyte border. ROI selection was performed without reference to either the eGFP or TH channels to minimise sampling bias. Paired ROIs were defined within each image to enable comparison of SC coverage across anatomical locations.

Fluorescence channels corresponding to eGFP and TH were separated and processed independently. To reduce background noise and improve signal continuity, images were subjected to spatial smoothing using the “Smooth” function. Thresholds were manually adjusted for each channel using the “Set Threshold” tool to ensure accurate segmentation of labelled structures. Binary masks were generated and refined using “Fill Holes” and “Watershed” functions to improve object continuity and separation. Schwann cell association with sympathetic axons was quantified by determining the proportion of TH-positive axonal signal that was associated with eGFP-positive SC processes within each ROI. Following thresholding, TH-positive signal coinciding with eGFP-positive signal was classified as SC-associated and expressed as a percentage of the total TH-positive signal. TH-positive signal lacking associated eGFP signal was classified as exposed axon. The proportion of exposed axon was calculated as the percentage of total TH-positive signal not associated with eGFP-positive SC processes. All image processing and analysis parameters were applied consistently across datasets. For each animal, 3-4 portal tract fields were analysed, and quantification was performed on tissue from n=3 animals.

#### Volume Correlative Light and Electron Microscopy (vCLEM)

Light microscopy was used to target volume sampling for ultrastructural analysis of peripheral nerve networks within the liver, as part of a hierarchical vCLEM workflow. This facilitated the targeting of SC positive regions, and permitted targeted, equitable sampling across animals to support unbiased quantification across comparable functional domains. Plp-directed eGFP fluorescence was used at three levels: first, to screen vibratome slices of liver to find SC positive slices (mm-cm scale), secondly to guide the trimming and collection of serial, ultrathin section arrays containing SC-associated regions (100’s μm scale); and thirdly, to refine the selection of ROIs within the array for high resolution serial section image acquisition (10’s μm scale).

Vibratome sections of paraformaldehyde/glutaraldehyde-fixed liver (left lobe; 150µm thickness, VT1200 S, Leica) from PLP-eGFP mice were incubated for 2 hours with Hoechst 33342 (1µm/ml) and tile-scanned using low-magnification confocal imaging (20x objective, non-immersion) to generate maps of the entire section and localise SCs. Additional structural context was obtained using vascular labelling (lectin) or cell-permeable fluorescent markers (including hydrazide or SiR-actin, 1µM), which enabled distinction of portal tracts from central veins and provided anatomical cues for consistent sampling. Tissue was not permeabilised to preserve membrane ultrastructure.

Sampling excluded the large hilar portal tracts (portal vein diameter >200µm) and the most distal terminal branches (<50µm). This approach enabled the generation of serial section arrays from anatomically comparable regions across animals while maintaining consistent tissue context. Following confocal mapping, vibratome sections were immediately processed in their entirety for volume electron microscopy (vEM) (see EM Sample Preparation, below).

Tissue containing SC positive regions of the portal tract (as described above) were excised from embedded sections under a dissecting microscope, using a razor blade, guided by matching vascular landmarks observed in the osmicated, resin embedded tissue to vasculature highlighted by lectin or hydrazide in confocal tile-scan images. Dissected tissue was mounted onto blank resin stubs using cyanoacrylate adhesive (superglue), and the block face trimmed for serial sectioning and collection onto ITO coated coverslips ^71^.

Serial sections were first imaged at low resolution (100nm pixel size) to capture the overall tissue architecture of the portal region. These datasets were cross-referenced with the fluorescence tile-scan images to guide the selection of ROIs for high-resolution acquisition for connectomic analysis. High-resolution acquisitions were performed at pixel sizes of either 5 or 10nm, depending on the experimental requirements.

Optimisation of pixel size was carried out to balance spatial resolution with total XY area coverage and the 10nm pixel size used for the majority of large-scale volume acquisitions.

### Electron Microscopy

#### Sample preparation for Transmission Electron Microscopy (TEM) and Array Tomography SEM (AT-SEM)

Array tomography scanning electron microscopy (AT-SEM) was used as the primary imaging modality for all vCLEM datasets and downstream connectomic analyses. Transmission electron microscopy (TEM) was used to validate ultrastructural features at higher resolution. Following fixation, samples for scanning electron microscopy were placed in glass vials and post-fixed in 1% osmium tetroxide /1.5% potassium ferricyanide for 60 minutes at room temperature in the dark, then washed in distilled water and dehydrated in a graded ethanol series (30%, 50%, 70%, 96% and 100%) with an intermediate step of 2 % uranyl acetate (UA) in 70% ethanol for 2.5 hours at 4°C. Resin Durcupan (Merck) was prepared by mixing Component A/M (48%), Component B (48%), Component C (2%) and Component D (2%). Samples were infiltrated with 50 % resin: 50 % propylene oxide (PO) solution O/N. Next morning, samples were put in 100 % resin for 24h. The following day, samples were placed in 0.2 mm thick Aclar film moulds (EMS) and baked at 60 °C for 48 hours.

#### Serial sectioning and AT SEM imaging

Resin embedded samples were trimmed using glass knives or razor blades in an ultramicrotome (UC6 or UC7 Leica Microsystems), to target portal tracts previously mapped by light microscopy. To facilitate the formation of continuous ribbons of serial sections and prevent section detachment during manipulation, a thin layer of adhesive was then applied to the leading edge of the final trimmed blockface^72^. Serial ultrathin sections (70nm) were cut using an Ultra Jumbo diamond knife (Diatome Ltd) to facilitate the organisation of numerous long ribbons on the large water surface, before being collected onto poly-l-lysine treated 22mm square indium tin oxide (ITO) coated glass coverslip, using the paperclip method^71^.

Sequential ribbons of ≤18mm in length were collected and aligned manually using eyelash manipulators and organised in ordered arrays. Once sufficient ribbons to fill a coverslip had been collected, sections were exposed to xylene/chloroform vapor to reduce compression artifacts, and the ribbons were docked to the coverslip (at air/water interface). The water was then withdrawn from the knife boat, allowing the section array to gently lie down onto the ITO coated glass coverslip. Sections were allowed to air dry, and subsequently post-stained with Reynolds’ lead citrate for 10 min to enhance contrast. Following extensive washes, sections were air-dried and stored until SEM imaging.

AT-SEM image acquisition was performed across multiple imaging sites using comparable SEM platforms (Zeiss Gemini series,) with Atlas 5’s 32kx 32k scan generator, and acquisition software (Fibics Inc.). While core acquisition parameters were standardised, minor differences in instrument configuration (e.g. stage bias, working distance, and probe current) and downstream processing workflows (including alignment strategies) were present between sites. These variations did not affect the identification or analysis of ultrastructural features but reflect site-specific optimisation of large-scale volume acquisition. Specific imaging parameters were as follows: N1: Gemini 300 FEG SEM with Sense backscatter detector, accelerating voltage 4.5kV, educed to a landing energy of 1.5kV by application of a stage bias of 3kV, and at a working distance of 5mm, 5nm x 5nm x 70nm voxels. N2: Gemini 460 FE-SEM with backscatter detector (BSD), accelerating voltage 6.5kV, reduced to a landing energy of 1.5kV by application of a stage bias of 5kV, and at a working distance of 3.5mm, 5nm x 5nm x 70nm voxels. N3-4: Gemini 450 FE-SEM with sense backscatter detector, accelerating voltage 2.0kV with current 300/500/300 pA (datasets D10/11/12 respectively), and at working distances of 6.8/5.8/5.9 mm (datasets D10/11/12 respectively), 10nm x 10nm x 70nm voxels. For very large-scale acquisitions (datasets D10-12), individual sections were imaged as multiple adjacent fields of view that were subsequently stitched to generate continuous section images. To save imaging time, hierarchical imaging was used to identify ROIs at lower resolution and permit customised high resolution acquisition geometries, to avoid imaging large empty regions (e.g. vascular lumens such as the PV). This strategy reduced acquisition time and enabled efficient scaling of large volumetric datasets for connectomic analysis.

#### TEM sectioning and imaging

Ribbons of 70nm thick sections were cut and 1-3 sections were collected per formvar coated slot grid. Sections were lead citrated for 7-10min, washed in H_2_O and dried for 20min. TEM imaging was done on a FEI Tecnai G2 Spirit Bio-Twin electron microscope and either a Morada G2 camera with iTEM software (Olympus Soft Imaging Solutions) or a Xarosa digital camera (EMSIS GmbH), controlled by Radius software (Version 2.1).

### Electron Microscopy Image processing and analysis

#### Registration of light and electron microscopy data and serial section alignment

Characteristic asymmetric patterns of blood vessels and nuclei were used as unbiased fiducials to register confocal slice images with EM images and confirm localisation of eGFP-positive SCs. Correlation of fluorescence images with electron micrographs was performed using Atlas 5 on the electron microscope or Adobe Photoshop or Microsoft Powerpoint for offline analysis. Serial image series or vEM datasets were aligned using Atlas 5 and/or TrakEM2^73^ and Fiji/ImageJ^69^. Where necessary, image alignments were refined in Webknossos using the AI alignment tool prior to annotation.

#### Annotation

To facilitate interactive online navigation, annotation, proof reading and collaborative team-based analysis of large 3D image volumes, aligned vEM datasets were uploaded and stored online in an open-access repository, Webknossos^31^. The full list of datasets and access to the vEM repository is available via links in Supplementary Table 1. In total, 12 datasets from 4 mice were analysed by a team of trained annotators, contributing >1000 hours of cumulative annotation time. All annotations were spatially linked to the underlying image data and stored as structured skeleton nodes and/or volumetric segmentations within Webknossos. Annotation workflows were validated through multi-annotator consensus; axon trajectories were independently verified by at least two annotators, while key features used for downstream analyses, including termini and varicosities, were confirmed by a minimum of three annotators. For volumetric features, including selected axons, varicosities and putative cellular and vascular targets of nerve contacts, objects were segmented within Webknossos using a combination of manual annotation and AI-assisted segmentation tools. The interpolation tool was applied where appropriate to bridge missing sections or mitigate segmentation or alignment artefacts, and to accelerate segmentation of large homogeneous objects such as nuclei. We thank the additional annotators: Raquel Laza Bulls, Carla Tobajas Gutierrez, Aurora Maria Pedro Bueno, Emma Mengual, Sergio Romero Rodriguez, Victor Llopis Maset, Emma Temprano Arenas, Ginevra Falco, Natalia Cormenzana, Jordi Xavier Lara Bejarano, Lucia Martinez Justicia, Nuria Bello Parea, Raquel Esteo Marco, Yara Vanherle, Inés Martinez Fernández, José Antonio del Saz, Anais Carabaño Llop, Claudia Ravello, Lidia Escuriet Cabo, Marta Miró Tudela, Olga Climent Requena, Nuo Yuan.

#### Axon trajectories

Axons were manually reconstructed using the “skeleton” tracing tool, generating continuous node-based 3D representations of individual nerve fibres across serial sections. Axon path length was measured using the tree length function.

#### Axon terminal analysis

Axon terminals were identified within vEM datasets based on the termination of reconstructed axon skeletons within the captured volume. End point node coordinates were exported to spreadsheet software (Google Sheets and Microsoft Excel) for consensus review by a team of three annotators and then downstream analysis (Table S2).

Terminals were defined as distal endpoints of axons that did not continue into further branching structures within the imaged volume. To distinguish axon terminals from large distal varicosities, only endpoints located at least 3µm from the nearest upstream branch point were classified as terminals. Distances were calculated along the axon skeleton trajectory in Webknossos using Path Length function. This criterion ensured large bulbous varicosities were not misclassified as terminal structures. All identifiable axon endpoints meeting these criteria within the analysed datasets, were included in downstream analysis. In a small number of cases, candidate axon termini were excluded from further analysis following consensus review by three annotators when sectioning artefacts, or incomplete volume capture prevented unambiguous ultrastructural interpretation.

Terminals were segmented and reconstructed in Webknossos using the Compute Mesh (ad-hoc) function to determine their 3D structure and were further classified according to morphology and local ultrastructural characteristics into distinct categories, including blunt and bulbous terminal morphologies. Terminal contacts with target cells were defined by continuous close membrane apposition between the axonal terminal membrane and the membrane of a neighbouring target cell across at least two consecutive serial sections without intervening cellular structures. For vascular and ductal structures, contacts were defined by close apposition of the axonal membrane to the associate basal lamina. A total of 54 axon terminals were identified and analysed across 11 reconstructed volume datasets from n=4 animals (datasets: D01_N1, D02_N1, D03_N1, D04_N1, D06_N2, D07_N2, D08_N2, D09_N2, D10_N3, D11_N3, D12_N4, Table S2).

For Figure 5H, the anatomical locations of all identified axon terminals were manually mapped onto a simplified schematic representation of the portal tract. Terminal positions were assigned based on their location relative to major portal tract landmarks, including the portal vein (PV), hepatic artery (HA), bile duct (BD), lymphatics (L), and surrounding hepatocyte layer, as determined from the corresponding vEM datasets. Because individual datasets sampled different regions of the portal tract, terminal locations from multiple datasets were combined to generate a composite overview of terminal distribution across the portal tract. The schematic therefore represents a summary of terminal locations observed across all analysed volumes (n=4 animals). Each terminal was colour-coded according to the identity of its associated target cell type. Terminals contacting multiple cell types were represented using the corresponding combination of colours. Terminals for which no associated target cell was present were classified as “no target”.

#### Axonal varicosity analysis

Axonal varicosities were identified along reconstructed axon skeletons within Webknossos. Varicosities were defined as axonal diameter ≥2-fold increase relative to the mean inter-varicosity axon diameter. To quantify varicosity-associated cellular interactions, reconstructed axons were sampled across multiple volume datasets from independent animals. To minimise sampling bias, axons were selected from different regions of the analysed portal tract volumes and from distinct anatomical contexts. Comparable cumulative axon lengths were analysed from each animal (1934.9µm, 1127.7µm, and 1571.2µm, respectively). All varicosities identified along the full reconstructed length of each sampled axon were included in subsequent analyses. Varicosities were further classified according to the anatomical context of the axon. Meshwork axons were defined as individual axons or small bundles, whereas bundle-associated axons were located within larger nerve bundles surrounding the hepatic artery and bile duct. Analyses of varicosity distribution and cellular contacts were performed separately for meshwork and bundle-associated axons.

Contacts between varicosities and target cells were defined using the same criteria applied for terminal contacts, based on continuous close membrane apposition across at least two consecutive serial sections without intervening cellular structures. A total of 673 individual varicosities in 102 axons were analysed from n=4 mice. Of these, 484 varicosities were identified from 60 meshwork axons (3.03 mm cumulative axon length, n=3 mice) from datasets: D01_N1, D02_N1, D03_N1, D06_N2, D10_N3 and 189 on 42 bundle-associated axons (1.84 mm cumulative axon length, n=4 mice) from datasets: D01_N1, D02_N1, D03_N1, D06_N2, D10_N3, D12_N4. In total, meshwork and bundle-associated varicosities formed 581 and 96 individual nerve-cell contacts, respectively (Table S2).

For Figure 6C, the anatomical locations of axonal varicosities were mapped onto a simplified schematic representation of the portal tract. 50 varicosities were randomly selected from the varicosity contact dataset (Table S2), comprising varicosities identified from n=3 animals. Varicosity positions were assigned based on their location relative to major portal tract landmarks, as for terminals above.

For Figure 7A, representative meshwork SC-free and -associated axons were selected from 3 animals to illustrate spatial distribution of varicosities along individual axon trajectories. Axons of similar length were chosen from each category for comparison, Varicosities were colour-coded according to identity of associated target cell type, varicosities lacking a cell contact were classified as “no target”, while varicosities associated with SCs were additionally indicated by a green outline.

#### Morphological identification and classification of cell types

Target cell types were identified directly within vEM datasets using established ultrastructural criteria, as summarised in Figure S4B. Cell classification was based on a combination of cellular and nuclear morphology, subcellular features, anatomical location, and the presence or absence of a basal lamina and a primary cilium ^74^. Hepatocytes were identified by their location abutting the portal tract, their polygonal morphology, centrally located round nuclei, abundant glycogen granules, numerous large mitochondria, and the presence of bile canaliculi. Cholangiocytes were identified as epithelial cells lining the biliary tree, including bile ducts and ductules, and were distinguished by their columnar morphology, apical specialisations (including microvilli and primary cilia), and continuous basal lamina. Identification of portal tract macrophages was based on their irregular morphology, presence of lysosomes, phagosomes, vacuoles and other electron-dense intracellular compartments characteristic of phagocytic cells. These cells lacked a basal lamina and a primary cilium, as confirmed by identification of a centriole pair adjacent to the Golgi apparatus during manual annotation^75^. Cells classified as fibroblasts within the portal tract were identified by their elongated spindle-shaped or reticular morphology, extensive rough endoplasmic reticulum, absence of basal lamina and presence of a primary cilium. Subclassification of macrophages and fibroblast populations was not performed for the purposes of this study.

#### Axonal diameter analysis

For Figure 2D-E, axonal diameter was quantified by averaging diameter measurements obtained from three axonal segments located between adjacent varicosities along each axon. For axons lacking varicosities, diameter was calculated by averaging at least three measurements taken along the axon trajectory. Measurements were performed using the distance tool, with 3D orientation of the axon taken into account to ensure accurate cross-sectional assessment. Diameter analysis was performed on 102 axons from n=3 mice, datasets: D01_N1, D02_N1, D03_N1, D10_N3, D12_N4.

#### Varicosity density analysis

For Figure 2D, distance between varicosities was calculated as 1/ (number of varicosities / unit axon length). All axons included in the axon varicosity analysis (n=4, datasets: D01_N1, D02_N1, D03_N1, D06_N2, D10_N3, D12_N4) were analysed. Axons were randomly sampled from nerve structures spanning a range of bundle sizes (1-50 fibres in cross section) and from multiple regions throughout the analysed volumes. Axon length was measured using the skeleton tree length function, and varicosity density was calculated as the total number of varicosities divided by the corresponding axon length. Varicosity density analysis was performed on 102 axons from n=4 mice.

#### Single axon location analysis

For Figure S2B, the anatomical location of single axons was manually mapped within each dataset. The percentage of single axons (>95% of its length is single) located at the hepatocyte border was calculated relative to the total number of single axons identified within that dataset. A total of 66 single axons from n=4 mice were analysed from datasets: D01_N1, D02_N1, D03_N1, D06_N2, D10_N3, D12_N4.

#### Spatial clustering analysis

For Figure S2K, clustering analysis was performed in MATLAB R2022b (MathWorks). Three volumetric datasets from 3 separate animals (datasets: D01_N1, D10_N3, D12_N4) were analysed independently and treated as biological replicates. Segmented varicosities were exported at a voxel resolution of 40 × 40 × 70 nm in x, y, and z respectively and imported as binary 3D image stacks (tif) and loaded using “tiffreadVolume”. Object centroids were extracted from the binary volumes using regionprops3 with the Centroid property. All spatial analyses were performed in nanometre units. To define the biologically relevant spatial domain for the analysis, the skeletons of all analysed axons were imported from .nml annotation files. These files contained manually traced axonal skeletons. The skeleton coordinates were used as a spatial reference to determine the position of detected varicosities relative to the underlying axons. The spatial extent of the axonal skeletons was then used to define the biologically meaningful volume within which random point simulations were generated for the clustering analysis.

For each dataset, spatial organisation of varicosities was quantified using nearest-neighbour distance (NND) analysis. Pairwise Euclidean distances between all centroids were calculated using “pdist2”, and the minimum non-self-distance was taken as the nearest-neighbour distance for each point. The mean NND of the observed distribution was then calculated. To determine whether the observed spatial organisation deviated from complete spatial randomness, a Monte Carlo randomisation approach was implemented. For each dataset, random point patterns containing the same number of objects as the observed data were generated within the bounding box defined by the spatial extent of the observed centroids. Random coordinates were sampled from a uniform distribution across the x, y, and z dimensions of this volume. Nearest-neighbour distances were calculated for each simulated configuration using the same procedure as for the observed data. This process was repeated 999 times to generate a null distribution of mean NND values expected under complete spatial randomness. Observed mean NND values were compared with the simulated null distribution to compute a z-score and Monte Carlo p-value, indicating whether varicosities were significantly clustered or dispersed relative to random spatial organisation. To visualise spatial organisation across biological replicates, the full distribution of nearest-neighbour distances was calculated for both observed and simulated point patterns for each dataset. Histograms were generated using a common set of bins across animals and normalised to probability density. Mean distributions across animals were then calculated and plotted together with the SEM.

#### SC coverage/exposure analysis of terminals

For Figure 3J, SC association at nerve terminals was defined based on direct membrane apposition observed in vEM datasets at the terminal. Axonal segments were classified as SC-associated, if any SC contact was made at the terminal, independent of the degree of SC coverage. Axon termini lacking such contact were classified as “free” (non-associated). In cases where SC-association status was ambiguous, classification was resolved by consensus between annotators or excluded from analysis. For SC-associated terminals, the degree of SC coverage was calculated as a percentage of axon perimeter within a 2D section. A minimum of three sections was analysed for each terminal, and the resulting measurements were averaged to obtain a single SC coverage value.

#### Analysis of axonal and varicosity exposure

For Figures 4E-4F, exposure of axons and varicosities was quantified from cross-sectional images using FIJI (ImageJ) and Webknossos. Exposure was defined as the proportion of the perimeter not in contact with SCs or neighbouring axons within the same bundle. Membrane regions contacting the extracellular matrix or other cell types, were classified as exposed.

For axonal analysis, individual axons were sampled throughout the reconstructed volumes by navigating to randomly selected regions within the dataset. Cross-sectional axon profiles were identified, and the total axonal circumference was measured using the Fiji freehand selection tool. The lengths of membrane contacting SCs or neighbouring axons (covered) or exposed to the surrounding tissue were measured manually using the freehand line tool. For each axon, the proportion of exposed membrane was calculated relative to the total axonal circumference. Axons were further classified according to bundle size as single axons (1 axon), small bundles (2–5 axons), or large bundles (>5 axons). A total of 245 axons from n=3 mice were analysed from datasets: D01_N1, D02_N1, D03_N1, D10_N3, D12_N4.

For varicosity analysis, cross-sectional views were obtained by sampling annotated varicosity centroids in WebWnossos. The total varicosity perimeter was measured using the distance measurement tool, and the lengths of membrane contacting SCs and neighbouring axons were measured separately. Varicosity exposure was subsequently calculated as proportions of the total varicosity perimeter using the same criteria as for axonal exposure above. A total of 393 varicosities from n=3 mice were analysed from datasets: D01_N1, D10_N3, D12_N4.

#### SC coverage of meshwork analysis

For Figure S3C, SC coverage was quantified for meshwork axons. Analysis was restricted to datasets in which all meshwork axons had been completely reconstructed and skeletonised. Each axon was examined along its entire length by sequential inspection of serial sections to identify regions with or without SC association. Meshwork axons were defined as individual axons or within bundles containing <10 axons. The path lengths of SC-associated and SC-free segments were measured using the Webknossos skeleton path length tool, and SC coverage was calculated as the percentage of the total reconstructed axon length covered by SCs. A total of 169 meshwork axons from n=3 mice were analysed from datasets: D01_N1, D02_N1, D03_N1, D06_N2, D10_N3.

#### Free axon location analysis

For Figure S3E, the anatomical location of SC-free axons was manually mapped within each dataset. The percentage of free axons located at the hepatocyte border was calculated relative to the total number of free axons identified within that dataset. A total of 64 free axons from n=4 mice were analysed from datasets: D01_N1, D02_N1, D03_N1, D06_N2, D10_N3, D12_N4.

#### SC association/exposure analysis along axons

For Figure S3K, axons with partial SC contact were identified within the volumes, the spatial organisation of SC association along axons was quantified by measuring the lengths of alternating SC-associated and SC-free regions along the length of an axon. Using tools on Webknossos, boundaries between SC-associated and -free regions were manually annotated along reconstructed axon skeletons. Consecutive boundary annotations defined individual axon lengths that were either associated with SCs (A) or exposed (B). Lengths were measured using the skeleton path length function. A total of 47 axons exhibiting intermittent SC-associated and exposed regions were identified across volume datasets from n=3 animals.

#### Distance to target cell analysis

For Figures S4C and S5E, the distance between an axon terminal/varicosity and a target cell was measured using Webknossos. All identified axon terminals were included in the analysis. For varicosities, ∼10 interactions per target cell type were randomly sampled from Table S1 from each animal, where available. Measurements were performed in a cross-sectional (XY) view by determining the shortest distance between the axonal membrane and the membrane of the adjacent target cell using the distance measurement tool. For each terminal/varicosity-target cell interaction, a minimum of three measurements were obtained at representative positions along the interface and averaged to generate a single distance value. Distance measurements were used to characterise the spatial relationship between terminals and target cells, ranging from near membrane apposition to interactions separated by a basal lamina. Given the lateral (XY) resolution of the AT-SEM datasets (5-10 nm per pixel), measurements approaching the lower range of observed distances should be interpreted as near-membrane apposition rather than definitive membrane contact.

#### Movies

Movies were generated from 3D light-sheet microscopy and vEM datasets. For Movie S1, animations were generated in Imaris (Oxford Instruments) by navigating through and rotating native light-sheet fluorescence data. For Movie S4 3D-rendering and movies of segmented vEM data were prepared using Amira^76^. Additional animations were generated directly in Webknossos^31^ using the built-in animation tool. Exported animations were assembled, annotated and edited using iMovie and Keynote software (Apple Inc.) prior to final export as MP4 files.

#### Statistical analysis

Statistical analysis was performed using Graphpad Prism software. Unless stated otherwise, data is presented as mean ± standard error of the mean (SEM) and is representative of at least 3 animals or independent cultures. Data were analysed by unpaired two-tailed t-test, one-way ANOVA, or two-way ANOVA, as appropriate, followed by multiple comparisons tests, when applicable, as indicated by the Figure Legends. In all cases, statistical significance was considered at p<0.05, and further indication of the p-value is indicated by (*): *p<0.05, **<0.01, ***<0.001, ****<0.0001, ns=not significant (p>0.05). In some cases, the recorded p-value is shown but not all comparisons are shown. The number of animals or biological repeats used in each experiment is stated in the Figure Legends.

